# Predictors Recover Most of the Metagenomic Signal for Antibiotic Resistance Gene Occurrence: A Cross-City Test of Geographic Transferability in Urban Wastewater

**DOI:** 10.64898/2026.09.24.753270

**Authors:** Somayeh Falahati Khanaman, Bichar Dip Shrestha Gurung, Shiva Aryal, Mengistu Geza Nisrani, Venkataramana Gadhamshetty, Etienne Z. Gnimpieba

## Abstract

Antibiotic resistance genes (ARGs) travel from cities into rivers and coastal waters through wastewater treatment plants. Monitoring them normally requires metagenomic sequencing, which is sufficiently costly that most utilities can sample only occasionally. Weather and location data are freely available on a daily basis for virtually any location worldwide, making them attractive predictors for identifying where limited sequencing resources should be prioritized. However, whether such models generalize to previously unseen cities remains largely untested, as most published studies train and evaluate models within the same catchments, thereby assessing interpolation rather than geographic transferability. To address this gap, we paired 235 wastewater metagenomes collected from five European cities with 23 abiotic predictors spanning geospatial, meteorological, hydrological, radiative, and temporal domains. Model performance was evaluated using leave-one-group-out (LOGO) cross-validation, in which all samples from one city were withheld for testing while the remaining cities were used for training. Cat-Boost achieved a median LOGO ROC–AUC of 0.929 and a median F1-score of 0.750. Using freely available environmental reanalysis predictors (meteorological, hydrological, radiative, geospatial, and temporal) — without any metagenomic sequencing — CatBoost achieved a mean ROC-AUC of 0.722, recovering 78% of the predictive performance of the full omics-integrated model. Removing latitude and longitude reduced ROC–AUC by only 0.003, whereas replacing random cross-validation with city-wise validation reduced ROC–AUC by 0.052. Predictive performance varied across ARG classes, ranging from a ROC–AUC of 0.981 for *β*-lactam resistance genes to 0.762 for glycopeptide resistance genes. These findings demonstrate that freely available environmental reanalysis predictors — spanning meteorological, hydrological, radiative, geospatial, and temporal domains — recover 78% of the predictive signal for antibiotic resistance gene occurrence in urban wastewater, allowing scarce sequencing capacity to be directed to the catchments where it changes a decision.

**Highlights:**

- Environmental features predict wastewater ARG risk without deep sequencing (ROC-AUC 0.722)
- CatBoost generalizes to unseen cities, reaching a LOGO ROC-AUC of 0.929
- Geographic transferability exceeds coordinate contribution (cost 0.052 vs 0.003)
- The framework prioritizes sequencing resources in data-poor regions at low cost

## 1. Introduction

The widespread global use and misuse of antibiotics in human healthcare, agriculture, aquaculture, and livestock sectors have emerged as major public health concerns with profound environmental implications[1, 2]. Antimicrobial resistance (AMR) is projected to cause approximately 10 million deaths annually by 2050, surpassing cancer as one of the leading causes of global mortality[3]. AMR can complicate the treatment of common bacterial infections, including pneumonia, bloodstream infections, urinary tract infections, and tuberculosis, when the causative pathogens harbor antibiotic resistance genes (ARGs) [4]. At the molecular level, ARGs, the genetic determinants underlying resistance mechanisms, are recognized as key drivers of this crisis because they can disseminate rapidly across environmental compartments through complex ecological networks. The environment serves as a major reservoir for ARGs, with continuous anthropogenic inputs promoting their accumulation in terrestrial and aquatic ecosystems and facilitating their transmission through environmental pathways, thereby accelerating the global spread of AMR. Recent studies have further highlighted the urgency of this issue, reporting that approximately 23.8% of identified ARGs in global environments pose potential human health risks because of their high transmissibility to pathogenic bacteria[5]. Consequently, ARGs are increasingly recognized as emerging environmental pollutants that require rapid, reliable, and scalable monitoring and mitigation strategies to limit their continued dissemination across ecosystems[6, 7].

Despite intensified surveillance efforts, existing ARG detection approaches, including culture-based techniques, molecular assays, metagenomic sequencing, and computational methods, continue to face significant limitations in terms of cost, scalability, and geographic applicability[8]. Shotgun metagenomic characterization of one wastewater sample requires DNA extraction, library preparation, sequencing to sufficient depth to recover low-abundance determinants, and an assembly and annotation pipeline. Turnaround is measured in days to weeks. The per-sample cost restricts most municipal programmes to occasional sampling campaigns rather than routine monitoring. Although each approach offers distinct advantages, none provides a comprehensive and cost-effective solution for large-scale environmental monitoring across diverse geographic regions. More importantly, most existing predictive frameworks rely primarily on sequence-based omics data while potentially overlooking the broader environmental context in which ARGs emerge, persist, and spread. The complex interactions among meteorological, hydrological, and geospatial factors that influence ARG dynamics remain insufficiently characterized, limiting the development of robust and geographically transferable surveillance models capable of operating independently of direct metagenomic measurements.

A second limitation concerns how such models are validated. Predictive studies of ARG dynamics have generally reported performance under random cross-validation, in which training and test partitions are drawn from the same sites and often the same sampling campaigns[9, 10, 11]. Wastewater samples from one catchment share their meteorological forcing, their contributing population, and their treatment configuration, so random partitioning places closely related observations on both sides of the split. The resulting estimates describe interpolation within a monitored system. A surveillance programme that deploys a model requires extrapolation to an unmonitored one. Whether the reported skill of environment-informed ARG models survives the removal of an entire city from training has not been tested. Wastewater treatment plants (WWTPs) represent critical convergence points for ARG accumulation and dissemination because they receive combined inputs from hospital effluents, industrial discharges, domestic wastewater, and agricultural runoff[12]. Although WWTPs employ multiple treatment processes, ARGs are not completely removed, resulting in their continuous release into receiving aquatic environments[13]. Environmental variables such as temperature, humidity, wind speed, precipitation, and geographic location have been shown to influence contaminant transport, microbial community composition, and ARG persistence in these systems[14, 15, 16, 17]. These abiotic factors affect bacterial survival, horizontal gene transfer, and the persistence of extracellular ARGs, highlighting the importance of integrating environmental predictors with omics-derived information to better characterize ARG dynamics in wastewater ecosystems[18, 19]. Despite growing evidence supporting these relationships, the systematic integration of meteorological, geospatial, temporal, and omics-derived predictors within a unified machine learning framework that generalizes across geographically distinct environments remains largely unexplored.

This study treats geographic transferability as the primary experimental variable rather than as a secondary robustness check. We paired 235 wastewater metagenomes from five European wastewater treatment plants (WWTPs) with 23 abiotic predictors and evaluated ARG occurrence prediction under Leave-One-Group-Out (LOGO) cross-validation, so that the reported performance estimates reflect transfer to an unmonitored catchment.

Three questions organize the analysis. First, how much predictive skill survives geographic transfer (Sections 4.3 and 4.4) Second, how much of that skill is carried by abiotic predictors that are freely and continuously available, rather than by metagenomic features that must be generated for each sample (Section 4.7). Third, is the remaining predictive skill attributable to transferable environment–resistome relationships or to spatial autocorrelation encoded in the geographic coordinates (Section 4.8). We address these questions by quantifying the performance changes associated with each experimental design choice and synthesize the findings into a Surveillance Deployment Readiness (SDR) framework (Section 4.11 and Table 8), which defines the conditions under which the proposed framework is suitable for guiding environmental surveillance and monitoring efforts.

## 2. Background

### 2.1. Environmental Sources and Dissemination Pathways of ARGs

An antibiotic resistance gene is a segment of bacterial DNA encoding proteins or enzymes that confer resistance to antibiotics through mechanisms such as enzymatic inactivation, target site modification, or efflux pump activation. Although ARGs occur naturally within bacterial populations, their prevalence has increased substantially due to selective pressure imposed by the widespread and often indiscriminate use of antibiotics [15, 20]

ARGs are disseminated through multiple environmental pathways originating from diverse anthropogenic sources.

Wastewater treatment plants are widely recognized as critical hotspots for ARG proliferation, where elevated concentrations of nutrients, residual antibiotics, heavy metals, and disinfectants create strong selective pressure that promotes ARG persistence and horizontal gene transfer among microbial communities [21, 22]. Although a range of treatment technologies, including biological methods such as membrane bioreactors and constructed wetlands, chemical approaches such as ozonation and chlorination, and physical methods such as UV irradiation and membrane filtration, have been deployed to reduce ARG loads, their removal efficiency remains incomplete and varies across treatment systems[13]. For example, a recent study of multiple wastewater treatment plants reported an average reduction of 88.4% in total ARG abundance, with removal efficiencies ranging from 63.2% to 94.2% among individual plants [23]. Consequently, residual ARGs and antibiotic-resistant bacteria can still be discharged into receiving water bodies.

Hospitals constitute another significant ARG source, driven by intensive antibiotic use and the high prevalence of resistant pathogens. Hospital waste streams, including urine, feces, and liquid effluents, frequently carry residual antibiotics and ARG-bearing microorganisms at concentrations and diversity levels markedly higher than those found in domestic wastewater [24]. The detection of ARGs in groundwater near urban hospitals further indicates that inadequately treated hospital effluents contribute to broader environmental contamination, with sludge disposal representing an additional dissemination route via horizontal gene transfer [25, 26].

Industrial wastewater represents a further major ARG introduction pathway, particularly in heavily industrialized regions where effluents contain elevated levels of antibiotics, heavy metals, and chemical contaminants that impose selective pressure on microbial communities [27]. The pharmaceutical manufacturing sector plays a particularly prominent role, as production effluents may harbor exceptionally high ARG concentrations, effectively functioning as environmental resistance hotspots [28]. Clinically relevant ARGs, including *bla*, *erm*, *tet*, and *sul*, have been detected in sediments and activated sludge associated with industrial wastewater, confirming their persistence through standard treatment processes [29, 30].

Agricultural activities constitute an equally important driver of ARG dissemination. A substantial proportion of administered antibiotics, including sulfonamides, tetracyclines, and fluoroquinolones, is excreted largely unmetabolized in animal manure, introducing both residual compounds and ARG-bearing bacteria into soils and water bodies [22]. In aquaculture systems, up to 90% of sampled aquatic bacteria have been reported resistant to at least one antibiotic [31]. Agricultural runoff further transports manure, biosolids, and ARG-laden soil particles into rivers and lakes, elevating ARG abundance and diversity in surface waters. Encouragingly, organic farming practices that restrict antibiotic use have been associated with approximately 31% lower ARG prevalence across livestock systems, underscoring the potential of management-level interventions to reduce environmental dissemination management-level interventions to reduce environmental dissemination [32, 33, 34].

Each of these source terms is coupled to the receiving WWTP through a transport process that depends on weather. Runoff mobilizes manure-associated and soil-associated ARGs into the sewer catchment during precipitation events. Atmospheric deposition delivers airborne resistance determinants at rates governed by wind and pressure fields. Evapotranspiration and soil moisture influence the dilution and residence time of the resulting load. Meteorological variables are therefore proxies for the transport terms linking a source to a plant, which provides the mechanistic basis for expecting abiotic predictors to carry recoverable signal in the absence of sequencing. This expectation is tested in Section 4.7.

### 2.2. Analytical Methods for Environmental ARG Surveillance

ARG detection methods are broadly categorized according to their underlying principles and scale of analysis, spanning traditional laboratory approaches to advanced computational frameworks.

Culture-based phenotypic methods assess resistance by evaluating microbial survival in the presence of antibiotics through techniques such as antibiotic susceptibility testing (AST), Kirby-Bauer disk diffusion, broth dilution, and agar dilution [35, 36, 37]. Although these approaches provide direct evidence of phenotypic resistance, they are fundamentally limited by the non-culturability of environmental microorganisms and their inability to reveal the specific genetic mechanisms underlying resistance.

Molecular genotypic methods address this limitation by targeting specific DNA markers and resistance genes within microbial genomes [36]. Techniques including PCR, quantitative PCR (qPCR), digital PCR (dPCR), loopmediated isothermal amplification (LAMP), and DNA microarrays enable sensitive and specific detection of known ARGs [38], though they confirm ARG presence without establishing active gene expression or functional resistance. Functional metagenomic methods offer a complementary approach by identifying ARGs on the basis of demonstrated biological activity rather than sequence similarity. Environmental DNA is extracted, fragmented, and cloned into a surrogate host, typically *Escherichia coli*, to construct metagenomic libraries that are subsequently screened for resistance phenotypes [39, 40]. A key advantage is the capacity to discover novel ARGs lacking homology to existing databases [41]; however, these methods are labor-intensive and constrained by heterologous expression requirements.

Computational sequence-based methods extend ARG detection to high-throughput sequencing datasets by comparing sequences against curated reference databases using alignment tools such as BLAST or DIAMOND [42, 43] or Hidden Markov Models (HMMs) [44]. Resources such as CARD [45] and ResFinder [46] enable rapid resistome profiling of complex environmental samples, though their effectiveness is inherently bounded by database completeness, limiting detection of novel or highly divergent ARG sequences.

Machine learning (ML)-based methods have recently emerged as powerful tools for ARG identification, classification, and abundance prediction. Classical algorithms, including Random Forest (RF), Support Vector Machines (SVM), eXtreme Gradient Boosting (XGBoost), and CatBoost [47, 48], alongside deep learning architectures such as Convolutional Neural Networks (CNNs) and Deep Neural Networks (DNNs) [49], have been widely applied to resistome analysis. Specialized frameworks such as DeepARG [50], ARGNet [9], and hybrid deep learning architectures [10] further illustrate the growing adoption of embedding-based representations for high-throughput resistome profiling.

Beyond sequence-based detection, ML approaches have been increasingly applied to predict ARG dynamics and identify key environmental and microbial drivers, effectively capturing nonlinear interactions and modeling complex relationships among environmental, microbial, and genomic features [5]. Lu et al. [11] applied multiple ML models to predict changes in ARGs and mobile genetic elements (MGEs) during biological nutrient removal, with CatBoost and RF demonstrating superior performance. Wang et al. [51] employed RF, XGBoost, and support vector regression (SVR) to model ARG dynamics during pig manure anaerobic digestion, with SVR achieving the highest predictive accuracy (*R*^2^ *≈* 0.78). Gradient boosted decision trees have also been applied to predict ARG abundance in microplastic-contaminated soils, identifying bacterial genera, climate variables, soil properties, and microplastic characteristics as primary drivers, with strong predictive performance (*R*^2^ *≈* 0.98) [52]. Collectively, these studies demonstrate the potential of ML to predict ARG abundance, uncover environmental determinants of resistance dissemination, and support AMR monitoring and management [53].

Each of these performance figures is conditioned on a validation design. The R² of about 0.98 reported for microplastic-contaminated soils[50] and the R² of about 0.78 for anaerobic digestion of swine manure[49] were both obtained under random partitioning of samples drawn from a common experimental system, and therefore describe interpolation within conditions the model has already seen. Table 6 makes this comparison explicit for the present work: the same CatBoost architecture on the same data returns different values depending on whether the partition is random or geographic. This literature establishes that environmental covariates carry ARG-relevant information within a system. Whether that information is portable between systems is the question addressed in Section 4.

### 2.3. Environmental Drivers of ARG Dynamics

Growing evidence demonstrates that abiotic and climatic conditions play fundamental roles in shaping the abundance, persistence, transport, and detectability of ARGs across diverse environmental compartments. Meteorological variables, particularly temperature and precipitation, strongly influence ARG dynamics by regulating microbial activity and facilitating the mobilization of antibiotic-resistant bacteria and genetic material across terrestrial, atmospheric, and aquatic environments [54]. Soils constitute one of the largest environmental reservoirs of ARGs and have been reported to contain up to 80% of clinically relevant antibiotics [55, 56, 17]. Soil physicochemical properties, including pH, moisture, salinity, and temperature, further regulate ARG persistence, with elevated temperatures generally associated with reduced ARG abundance [57, 58, 59]. Conversely, rainfall events can increase both the abundance and diversity of ARGs by transporting antibiotic-resistant bacteria and extracellular genetic material into soils and receiving surface waters [60].

In aquatic and coastal environments, meteorological and hydrological conditions similarly exert strong controls on ARG distribution and persistence. Temperature, precipitation, and water movement influence microbial activity, contaminant transport, and molecular detection outcomes, making them important predictors of ARG abundance [61, 62]. Physicochemical characteristics, particularly salinity, have also been associated with shifts in ARG composition, with elevated salinity linked to increased ARG abundance in regions of the Pacific Ocean [20]. At broader spatial scales, atmospheric processes further influence ARG dissemination, as higher wind speeds and diverse air mass trajectories transport distinct microbial communities and environmental DNA across geographic regions [63].

Large-scale climatic gradients also contribute to the global distribution of ARGs. Elevated ARG abundance has frequently been observed in high-latitude cold and boreal regions, where climatic seasonality and mobile genetic elements strongly influence resistance dissemination. Hydrological conditions regulate water-mediated transport of ARGs and microbial communities, whereas evapotranspiration, which integrates temperature, solar radiation, and land-surface moisture, serves as an important indicator of ecosystem water balance and moisture dynamics [64]. Likewise, atmospheric pressure and associated weather systems influence aerosol transport, humidity, and deposition processes, thereby affecting the long-range dispersal of ARG-carrying microorganisms [65].

Collectively, these findings confirm that atmospheric, hydrological, and climatic drivers govern ARG persistence, spatial movement, and detectability across environmental systems. This body of evidence forms the scientific rationale for integrating meteorological, geospatial, and temporal abiotic predictors alongside omics-derived features within ML-based ARG surveillance frameworks, the approach adopted in the present study. These mechanisms also generate a testable expectation for the interpretation in Section 4.6. If the model exploits the transport processes described above, the predictors it weights most heavily should be those that proxy transport and persistence: precipitation and soil moisture for runoff-mediated delivery, wind speed and surface pressure for aerosol-mediated delivery, and evapotranspiration for the water balance governing dilution. The weighting should also shift between maritime and continental cities as the dominant transport pathway shifts. If instead the model weights predictors that only index location, the mechanism is absent and the skill reflects spatial autocorrelation. Sections 4.6 and 4.8 evaluate these alternatives. Importantly, prior studies have demonstrated that environmental and climatic variables alone can achieve substantial predictive performance for ARG abundance and occurrence, even in the absence of direct metagenomic profiling [52, 62], providing empirical precedent for the abiotic-only evaluation conducted in Section 4.7.

Collectively, these findings confirm that atmospheric, hydrological, and climatic drivers govern ARG persistence, spatial movement, and detectability across environmental systems. This body of evidence forms the scientific rationale for integrating meteorological, geospatial, and temporal abiotic predictors alongside omics-derived features within ML-based ARG surveillance frameworks, the approach adopted in the present study. These mechanisms also generate a testable expectation for the interpretation in Section 4.6. If the model exploits the transport processes described above, the predictors it weights most heavily should be those that proxy transport and persistence: precipitation and soil moisture for runoff-mediated delivery, wind speed and surface pressure for aerosol-mediated delivery, and evapotranspiration for the water balance governing dilution. The weighting should also shift between maritime and continental cities as the dominant transport pathway shifts. If instead the model weights predictors that only index location, the mechanism is absent and the skill reflects spatial autocorrelation. Sections 4.6 and 4.8 evaluate these alternatives. Importantly, prior studies have demonstrated that environmental and climatic variables alone can achieve substantial predictive performance for ARG abundance and occurrence, even in the absence of direct metagenomic profiling [52, 62], providing empirical precedent for the abiotic-only evaluation conducted in Section 4.7.

## 3. Methodology

### 3.1. Computational Framework

The proposed computational framework integrates environmental and metagenomic data with machine learning to predict and interpret antibiotic resistance gene occurrence in urban wastewater. The complete seven-stage workflow is presented in Figure 1.

**Figure 1:**
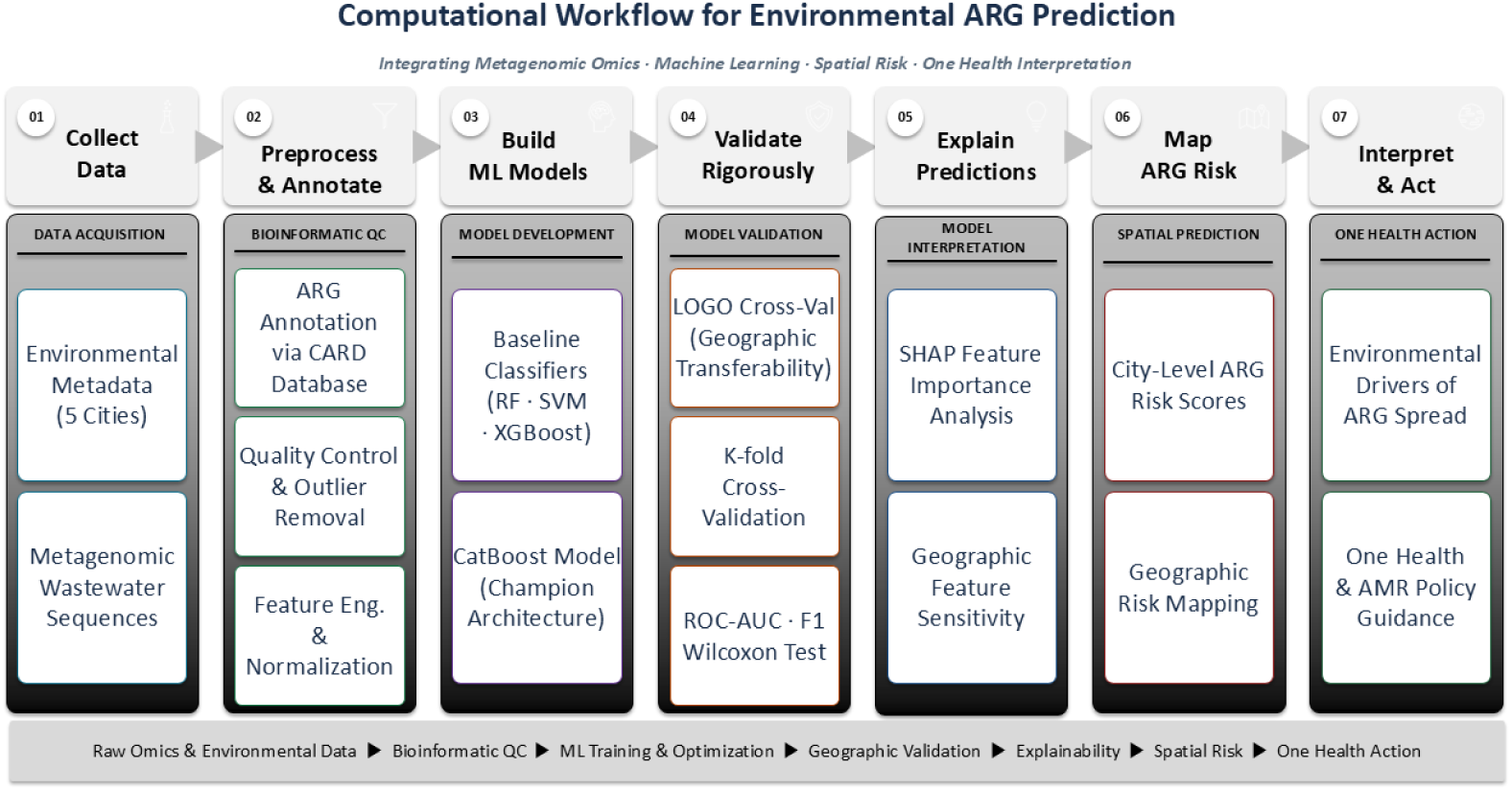
Computational workflow for environmental antibiotic resistance gene (ARG) prediction in urban wastewater, integrating metagenomic omics, machine learning, spatial risk, and One Health interpretation. The pipeline comprises seven sequential stages: (1) Data Acquisition — environmental metadata and metagenomic wastewater sequences were collected from treatment plant influents across five cities; (2) Bioinformatic QC — raw sequences were annotated against the Comprehensive Antibiotic Resistance Database (CARD), followed by quality control and outlier removal, then feature engineering and normalization of abiotic and omics-derived predictors; (3) Model Development — baseline supervised classifiers (Random Forest, Support Vector Machine, and XGBoost) were trained to predict binary ARG class occurrence, with a CatBoost model retained as the champion architecture; (4) Model Validation — geographic transferability was assessed by Leave-One-Group-Out (LOGO) cross-validation alongside K-fold internal cross-validation, with performance evaluated by ROC-AUC, F1-score, and pairwise Wilcoxon signed-rank tests; (5) Model Interpretation — SHapley Additive exPlanations (SHAP) feature importance and geographic feature sensitivity analyses quantified the relative contribution of abiotic, temporal, and microbial community predictors; (6) Spatial Prediction — citylevel ARG risk scores were derived from model outputs and rendered as geographic risk maps identifying high-burden urban areas; and (7) One Health Action — dominant environmental drivers of ARG spread were identified and translated into One Health and AMR policy guidance for surveillance and public health. CARD, Comprehensive Antibiotic Resistance Database; LOGO, Leave-One-Group-Out; ROC-AUC, Receiver Operating Characteristic — Area Under the Curve; SHAP, SHapley Additive exPlanations; ARG, antibiotic resistance gene; AMR, antimicrobial resistance.

#### 3.1.1. Prediction Target and Data Provenance

All models perform binary classification of ARG occurrence for each of the 15 resistance classes. Metagenomic sequence data derive from a single published source [66] (ENA BioProject PRJEB68319); no sequencing was performed for this study. Abiotic predictors derive from the NASA POWER reanalysis product, queried at each plant’s coordinates for each sampling date. Both sources are open and independently retrievable, and the analysis code is deposited (Data Availability).

### 3.2. Metagenomic Sequence Processing

Raw paired-end metagenomic sequencing data were obtained from the European Nucleotide Archive (ENA) under BioProject accession PRJEB68319, comprising longitudinal wastewater metagenomes collected from five European cities [66]. For computational feasibility, paired-end reads were processed from the subsampled FASTQ files used in this study. Raw read quality was assessed using FastQC and summarized with MultiQC [67]. Adapter trimming and quality filtering were performed using *fastp* [68] with pairedend adapter detection enabled, a minimum qualified Phred score of 20, a minimum retained read length of 50 bp, and two processing threads. No host-read depletion step was performed. High-quality reads were assembled into contigs using MEGAHIT [69], and protein-coding genes were predicted from assembled contigs using Prodigal in metagenomic mode, generating translated protein sequences in FASTA amino acid (.faa) format. Predicted protein sequences were analyzed using the Resistance Gene Identifier (RGI version 6.0.8) with DIAMOND alignment against the Comprehensive Antibiotic Resistance Database (CARD version 4.0.1; accessed May 2025) [45]. RGI was executed using protein sequence input with DIAMOND as the alignment tool and default detection criteria, retaining only Perfect and Strict ARG matches. For each sample, the number of predicted protein sequences assigned to each ARG (Best_Hit_ARO) was quantified from the filtered RGI outputs to construct an ARG-by-sample count matrix comprising 554 unique ARGs. No additional read-level normalization was applied during ARG identification; subsequent log-transformation and preprocessing of ARG count profiles are described in Section 3.4. ARG identifiers were mapped to CARD ontology annotation files (aro_index.tsv, aro_categories_index.tsv, and card.json) to standardize gene nomenclature and classify the detected ARGs into 15 antibiotic resistance classes. Of the 278 samples available in the original dataset, 235 samples with complete ARG profiles and corresponding environmental metadata were retained for downstream machine-learning analyses.

Two pipeline choices bound the interpretation of the results. First, assembly-based recovery detects ARGs only from contigs that assemble, and assembly efficiency scales with coverage, so a low-abundance determinant that fails to assemble at the subsampled depth is recorded as absent rather than as missing. The occurrence label for a rare class therefore depends partly on sequencing depth in that sample. Second, retaining only Perfect and Strict RGI matches excludes divergent environmental homologues by construction, so the resistome characterized here is the clinically annotated resistome rather than the total resistome.

### 3.3. Site Description and Data Collection

This study analyzed 235 publicly available wastewater metagenomes collected between 2019 and 2021 from WWTPs in five European cities: Bologna (Italy), Budapest (Hungary), Copenhagen (Denmark), Rome (Italy), and Rotterdam (the Netherlands). The dataset was originally generated and described by by Becsei et al. [66]. Raw paired-end sequencing data were downloaded from the European Nucleotide Archive (ENA) under project accession PRJEB68319 (Figure 2) .Together, these cities represent a diverse latitudinal and climatic transect, ranging from the oceanic conditions of Copenhagen and Rotterdam to the Mediterranean and humid subtropical environments of Rome and Bologna, with Budapest providing a continental inland contrast enabling assessment of ARG dynamics across markedly different environmental settings.

**Figure 2:**
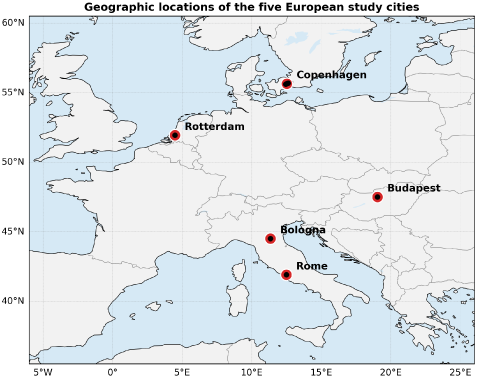
Geographic locations of the five European study cities.

The 2019-2021 sampling window spans the COVID-19 pandemic, during which antibiotic prescribing, hospital occupancy, and population mobility changed substantially and unevenly across these five countries. Sample distributions by year and city are reported in the Table S1.

Beyond climate, the five cities differ substantially in their anthropogenic profiles, which may further influence local ARG dissemination patterns. Northern and central hubs such as Copenhagen and Rotterdam are characterized by biotechnology, pharmaceutical manufacturing, and advanced agri-food logistics, whereas Bologna and Rome reflect a southern European profile dominated by machinery production, food processing, and tourism. Agricultural practices follow a similar north-to-south gradient, transitioning from precision, high-tech agri-logistics in northern cities to more intensive crop and livestock production in the south. Budapest occupies an intermediate position, combining grain and livestock agriculture with a diversified manufacturing and services economy. These contrasting economic, agricultural, and industrial profiles create distinct environmental signatures that potentially shape the local distribution and dissemination of ARGs (Table 1).

**Table 1:** Key features of the studied cities.

| City | Loc. | Climate | Agri. | Industry | Source |
| --- | --- | --- | --- | --- | --- |
| Bologna | North | Humid subtrop. | Crops, dairy | Machinery, food | [70] |
| Budapest | Central | Continental | Grain, livestock | Services, manuf. | [71] |
| Copenhagen | North | Mild oceanic | Agri-tech hub | Pharma, biotech | [72] |
| Rome | Central | Mediterranean | Crops, livestock | Tourism, gov. | [73] |
| Rotterdam | North | Maritime | Agri-food log. | Port, energy | [74] |

### 3.4. Abiotic Predictors and Omics Data

The study dataset comprised multidimensional abiotic predictors and high-throughput metagenomic data, which were integrated for subsequent machine learning analyses. A total of 23 abiotic predictors were assembled and categorized into five thematic domains: geospatial, meteorological, hydrological, radiative, and temporal.

Geospatial predictors included (LATITUDE) and (LONGITUDE). Meteorological predictors comprised near-surface air temperature (T2M), maximum and minimum air temperature (T2M_MAX and T2M_MIN), surface skin temperature (TS), dew point temperature (T2MDEW), wet-bulb temperature (T2MWET), relative humidity (RH2M), specific humidity (QV2M), wind speed (WS2M), and surface pressure (PS). Hydrological predictors included corrected total precipitation (PRECTOTCORR), surface soil moisture (GWETTOP), root-zone soil moisture (GWETROOT), profile soil moisture (GWETPROF), and evapotranspiration (EVPTRNS). Radiative predictors comprised all-sky downwelling shortwave radiation (ALLSKY_SFC_SW_DWN), all-sky downwelling longwave radiation (ALLSKY_SFC_LW_DWN), cloud fraction (CLOUD_AMT), and precipitable water vapor (PW). Temporal predictors consisted of relative week and weekof-year indicators used to capture seasonal variability.

Several predictors are near-deterministic functions of one another. T2M, T2M_MAX, T2M_MIN, TS, T2MDEW and T2MWET are six representations of the same thermal state, and RH2M, QV2M and PW are three representations of the same moisture state. Tree ensembles are robust to collinearity in that predictive performance is not degraded, but SHAP attribution is not: when two predictors carry the same information, the Shapley value is divided between them according to tree structure rather than mechanism. The SHAP rankings in Section 4.6 are therefore reported at the level of thermal, moisture and atmospheric-transport blocks rather than individual variables. The predictor correlation matrix is provided in the Figure S1.

The abiotic predictors exhibited clear spatial variation across the five study cities, providing a diverse environmental gradient suitable for subsequent machine learning analyses (Figure 3a).

**Figure 3:**
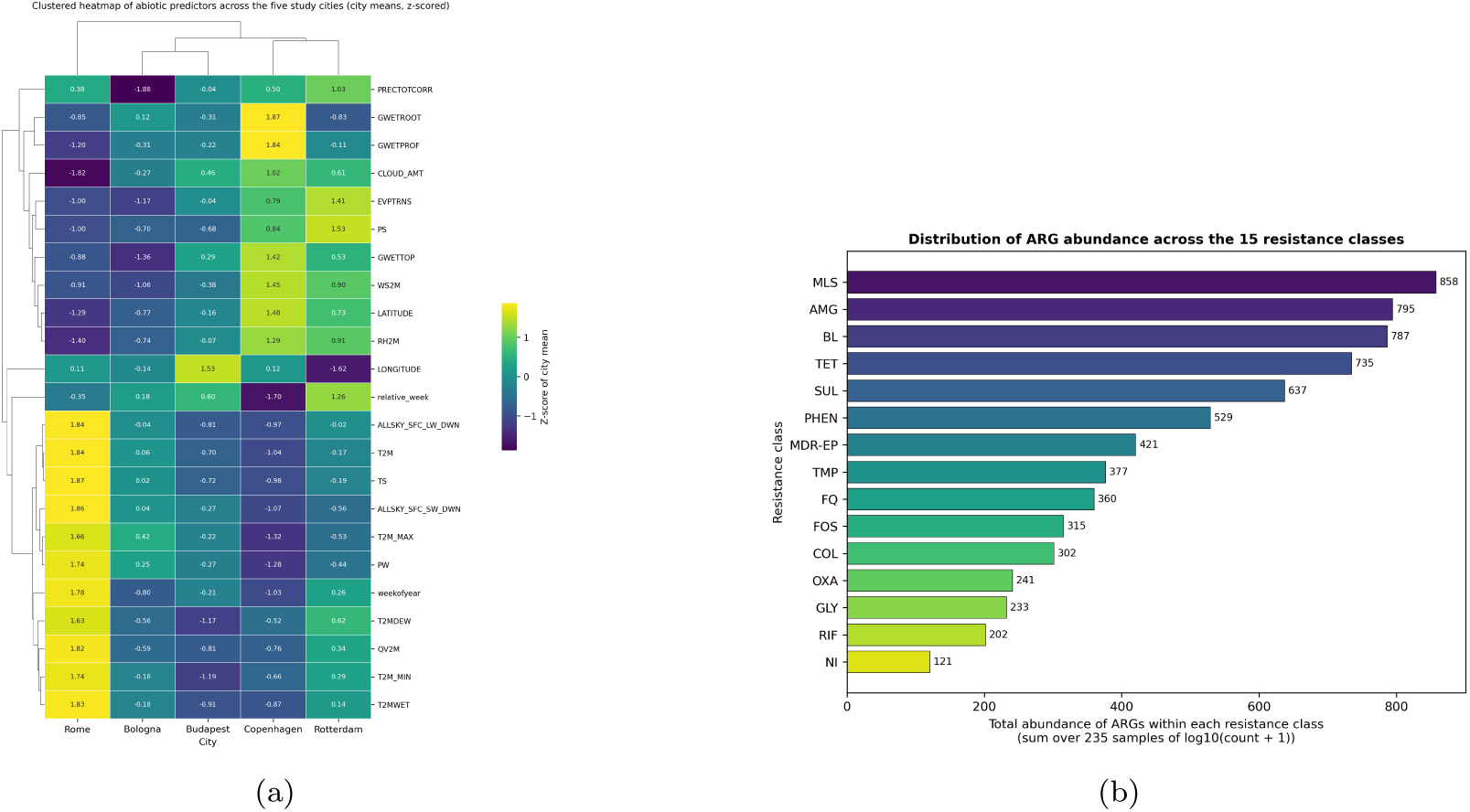
Characterization of abiotic predictors and ARG profiles across the five study cities. (a) Clustered heatmap of abiotic predictors showing distinct environmental gradients among the study cities. (b) Distribution of raw ARG abundance across the 15 resistance classes, calculated by summing the abundances of ARGs assigned to each resistance category across the five study cities.

The metagenomic dataset contained 554 unique ARGs detected across the five study cities. These genes were subsequently processed and grouped into 15 major antibiotic resistance classes during the bioinformatics preprocessing stage. (Figure 3b) provides an overview of the distribution of these ARG classes. Together, the abiotic predictors and metagenomic observations constituted the integrated dataset used for subsequent preprocessing and machine learning analyses.

### 3.5. Data Preprocessing

#### 3.5.1. Bioinformatic Layer: ARG Abundance Processing

A total of 554 unique ARGs were detected across the 235 wastewater samples collected from the five study cities. The input ARG matrix consisted of per-sample ARG counts derived from the filtered Resistance Gene Identifier (RGI) outputs, where each value represented the number of predicted protein sequences assigned to a given ARG (Best_Hit_ARO) within an individual sample.

Prior to metagenomic assembly, sequencing depth was standardized by randomly subsampling each paired-end sequencing library to 1,000,000 read pairs using seqtk sample. A fixed random seed (seed = 42) was applied consistently to both forward and reverse read files to maintain paired-end synchronization and ensure reproducible subsampling. Standardizing the input sequencing depth reduced the potential influence of unequal sequencing effort on downstream assembly, gene prediction, and ARG detection while providing a computationally tractable dataset for metagenomic analysis.

Because sequencing depth had already been standardized before assembly, no additional read-level normalization (e.g., reads per million mapped reads (RPM) or reads per kilobase million (RPKM)) was applied prior to model development. Raw ARG counts were inspected for missing values and zero observations before applying a base-10 logarithmic transformation to reduce distributional skewness while retaining zero values:

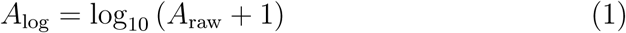

The log-transformed ARG matrix subsequently served as the omics-derived feature matrix for all supervised machine-learning models.

For ecological community analyses, the standardized ARG count matrix was converted to sample-wise relative abundances prior to calculating Bray–Curtis dissimilarity. Principal Coordinates Analysis (PCoA) was then performed to visualize compositional variation in reduced-dimensional space (Figure 4). Permutational Multivariate Analysis of Variance (PERMANOVA) indicated significant differences in resistome profiles among cities (*F* = 68.59*, R*^2^ = 0.54*, p* = 0.001). A test of multivariate homogeneity of group dispersions (PERMDISP) is also reported alongside the PERMANOVA results in the Table S2. These findings supported the use of LOGO cross-validation to evaluate model generalizability across geographically distinct urban environments.

**Figure 4:**
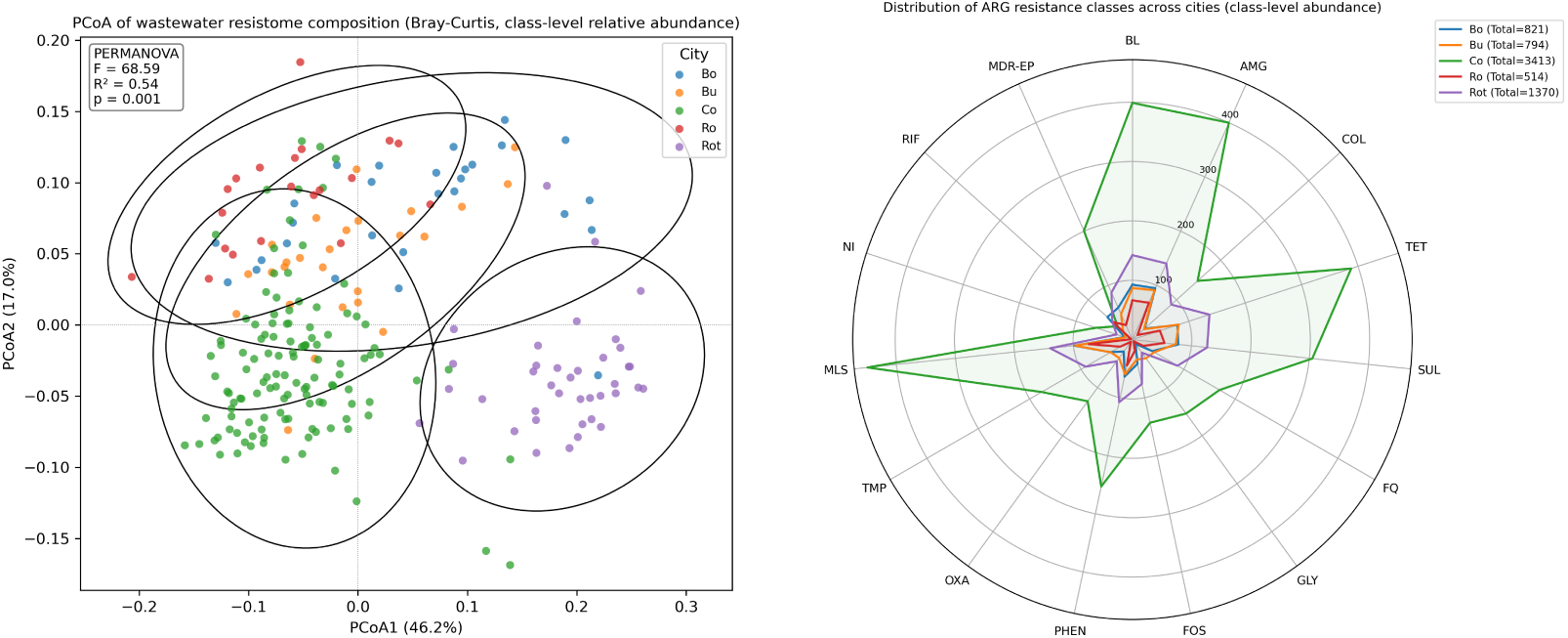
(a) Principal Coordinates Analysis (PCoA) of wastewater resistome composition based on Bray–Curtis dissimilarity calculated from sample-wise relative ARG abundances. (b) Radar plot showing the distribution of ARG resistance classes across the five study cities (Bologna (Bo), Budapest (Bu), Copenhagen (Co), Rome (Ro), and Rotterdam (Rot)) based on class-level abundances.

Following resistome characterization, individual ARGs were grouped into resistance classes using established classifications derived from the CARD and the Antibiotic Resistance Ontology (ARO) [45]. This yielded 15 major resistance classes: aminoglycosides (AMG), *β*-lactams (BL), tetracyclines (TET), sulfonamides (SUL), macrolide–lincosamide–streptogramin (MLS), trimethoprim (TMP), fluoroquinolones (FQ), phenicols (PHEN), glycopeptides (GLY), fosfomycin (FOS), colistin (COL), nitroimidazoles (NI), oxazolidinones (OXA), rifamycins (RIF), and multidrug/efflux pump-associated genes (MDR-EP) [45, 5]. Class-level abundance profiles were subsequently generated from the ARG abundance matrix and used as omics-derived predictor variables for subsequent machine-learning analyses. Figure 4 provides an overview of the distribution of these resistance classes across the five study cities.

In addition to resistance-class grouping, ARGs were categorized according to their reported environmental sources based on published ARG source annotations. Relative source contributions were subsequently calculated for exploratory characterization of the detected resistome.

#### 3.5.2. Environmental Layer: Abiotic Predictor Processing

Abiotic predictors were matched to each wastewater sample according to its sampling date and geographic location to ensure consistency between environmental and metagenomic observations. Prior to model development, the environmental dataset was inspected for missing values and potential outliers. Potential outliers were identified using the Interquartile Range (IQR) method as part of the data quality assessment. No observations were removed or modified based on the IQR criterion, as extreme environmental values (e.g., precipitation events) may represent genuine environmental conditions rather than measurement errors. All observations were therefore retained for subsequent analyses.

Continuous abiotic predictors were standardized using the StandardScaler transformation, which centers each variable to zero mean and scales it to unit variance. Standardization was required for Logistic Regression to ensure stable coefficient estimation and numerical optimization. Although feature scaling is not required for the tree-based models (Random Forest, XGBoost, and CatBoost), the same standardized predictor matrix was used across all models to maintain a consistent preprocessing pipeline. For both 5-fold crossvalidation and LOGO cross-validation, the scaler was fitted exclusively on the training data within each fold and subsequently applied to the corresponding validation data, thereby preventing information leakage from the held-out samples.

Together, the bioinformatic and environmental preprocessing layers preserved the biological characteristics of the ARG abundance data while generating a standardized and internally consistent feature set suitable for subsequent machine learning modeling.

### 3.6. Machine Learning Model Development and Validation

Four supervised machine learning algorithms were evaluated to predict ARG occurrence using thirty-eight integrated abiotic and omics-derived predictors: Logistic Regression(LR), RF, XGBoost, and CatBoost. LR, RF, and XGBoost were included as benchmark models because of their widespread application in environmental microbiology and antimicrobial resistance studies, providing well-established baselines for comparison.

CatBoost was selected as the primary classification model owing to its strong performance on heterogeneous tabular datasets and its ability to capture complex nonlinear relationships among environmental and metagenomic predictors [75]. As a gradient-boosted decision tree algorithm, CatBoost incorporates ordered boosting and symmetric tree construction, reducing prediction bias while improving generalization and computational efficiency. These characteristics make it well suited for modeling high-dimensional environmental datasets with complex interactions among predictors [76].

Baseline models—Logistic Regression, Random Forest, and XGBoost— were tuned using randomized hyperparameter search (RandomizedSearchCV, 30 iterations, optimized using ROC–AUC) to provide a fair comparison against the manually optimized CatBoost model. For each LOGO iteration, hyperparameter optimization was performed using an inner group-wise cross-validation scheme (GroupKFold) restricted to the training cities only, thereby ensuring that the held-out city’s data did not influence model selection at any stage. Logistic Regression was tuned over the inverse regularization strength (*C*), while using class-balanced weighting and the lbfgs solver. Random Forest was tuned over the number of trees, maximum tree depth, minimum samples per leaf, and the number of features considered at each split, with class-balanced weighting. XGBoost was tuned over the number of boosting iterations, maximum tree depth, learning rate, subsampling ratio, column subsampling ratio, and L2 regularization strength (*λ*). To account for class imbalance, the scale_pos_weight parameter was calculated separately within each training fold based exclusively on the training data, thereby preventing information leakage from the held-out city. All models were initialized using a fixed random seed (42) to ensure reproducibility. The fixed CatBoost model and the optimized baseline models were subsequently applied to the held-out city in each LOGO iteration, and predictive performance was evaluated across all five validation folds. The corresponding hyperparameter search spaces and final model settings are presented in Table S3.

To mitigate the effects of class imbalance, class-balancing strategies were applied consistently across all machine-learning algorithms. CatBoost used auto_class_weights=’Balanced’, which automatically adjusts class weights according to the distribution of positive and negative samples within each binary classification task. Logistic Regression and Random Forest employed class_weight=’balanced’, whereas XGBoost used a fold-specific scale_pos_weight calculated exclusively from the training data within each LOGO iteration. These approaches reduced bias toward the majority class while preventing information leakage from the held-out city.

Model generalizability was evaluated using a LOGO cross-validation strategy, in which all samples from one city were withheld as an independent test set while samples from the remaining four cities were used for model training. This process was repeated until each city had served once as the test set. Compared with conventional random cross-validation, LOGO prevents information leakage among geographically related samples and provides a more realistic assessment of model performance in previously unseen urban environments [77].

With five study cities, LOGO yields five validation folds. Consequently, summary performance statistics (e.g., mean, median, and standard deviation of ROC–AUC and F1-score) are derived from five geographically independent evaluations and should therefore be interpreted with appropriate caution. Furthermore, the validation folds are not exchangeable because each heldout city represents a distinct geographic and environmental setting; for example, predicting ARG occurrence in Mediterranean Rome constitutes a fundamentally different transfer task from predicting it in maritime Rotterdam. Consequently, the average performance across folds summarizes geographic transferability across these five urban environments rather than providing an estimate that can be directly generalized to all wastewater systems. To facilitate interpretation of this variability, per-city and per-class performance metrics are reported alongside the overall summary statistics (Table S4).

### 3.7. Model Evaluation

Model performance was evaluated using the F1-score and the area under the receiver operating characteristic curve (ROC–AUC), both computed according to their standard definitions. Mean, median, and standard deviation were calculated across the LOGO validation folds to summarize overall predictive performance, typical model behavior, and variability among geographically independent test sets.

The F1-score was selected because it provides a balanced assessment of precision and recall, making it particularly suitable for evaluating models under class imbalance. ROC–AUC was used as a threshold-independent measure of the model’s ability to discriminate between ARG occurrence classes across different decision thresholds. Because ROC–AUC may overestimate predictive performance under class imbalance, particularly when the negative class substantially exceeds the positive class, the area under the precision–recall curve (PR–AUC) was additionally reported as a complementary evaluation metric. Since positive predictions directly determine sequencing effort, precision provides a more informative measure of practical performance. Class-specific PR–AUC results for all 15 ARG resistance classes are provided in Table S4.

The mean performance represents the expected predictive capability across geographically distinct validation folds, whereas the median provides a robust estimate of the typical predictive performance that is less sensitive to extreme values arising from individual cities. Because LOGO validation comprised only five geographically independent test folds, both mean and median values are reported to provide complementary summaries of model performance across heterogeneous urban environments. The standard deviation was used to assess model stability and consistency across geographically independent validation sets, thereby quantifying the robustness of model generalization under varying environmental conditions. To statistically compare the predictive performance of the evaluated machine learning models, pairwise Wilcoxon signed-rank tests were performed using paired ROC–AUC, PR–AUC, and F1-score values obtained for each Target_Gene *×* held-out city combination under the LOGO cross-validation framework. Each paired observation therefore represented the performance of two competing models evaluated on the same ARG resistance class and the same held-out city, yielding 65 paired observations for ROC–AUC (following exclusion of folds with undefined values due to single-class test sets) and 75 paired observations for PR–AUC and F1-score. Because ARG resistance classes evaluated within the same held-out city share the same training/test partition and environmental covariates, these paired observations have a nested structure. Accordingly, the Wilcoxon signed-rank test was used as an exploratory non-parametric comparison of paired model performance, and inferential results were interpreted together with the reported effect sizes and descriptive performance statistics. To account for the three pairwise model comparisons performed for each evaluation metric, Holm’s sequential correction was applied to the resulting *p*-values. Statistical significance was assessed using an adjusted significance level of *α* = 0.05.

### 3.8. Model Interpretation

To improve model interpretability, SHAP were employed to quantify the contribution of individual predictors to model outputs [78]. Based on cooperative game theory, SHAP assigns an importance value to each feature for every individual prediction, allowing the contribution of both abiotic and omics-derived predictors to be quantified.

Global SHAP analysis was used to identify the overall importance of predictors across the complete dataset, whereas local SHAP values were employed to explain feature contributions for individual wastewater samples. SHAP decomposes a model’s output rather than the underlying environmental system. It identifies which predictors a fitted model relied upon, conditional on the correlation structure of the training data. Because several predictors exhibit substantial correlation, SHAP values should be interpreted as explanations of the model’s predictions rather than as evidence of causal drivers of ARG occurrence. Consequently, variables acting as proxies for unmeasured environmental processes may also receive high Shapley values. To assess model interpretability under geographically transferable conditions, all SHAP values presented in Section 4.6 were computed exclusively for the held-out test fold in each LOGO cross-validation iteration.

## 4. Results and Discussion

### 4.1. Ecological Reservoirs of Detected ARG Classes

The 15 ARG classes identified in this study have been previously reported across diverse ecological reservoirs in the literature, including hospitals, municipal wastewater, livestock production systems, agricultural environments, and natural ecosystems (Table 2). These literature-based associations are presented as interpretive context for the detected resistome rather than as source-tracking evidence from the sampled wastewater itself, since sample-level provenance was not directly measured in this study. Nonetheless, the fact that the detected ARG classes are, according to prior surveillance studies, commonly distributed among interconnected human, animal, and environmental reservoirs is consistent with the One Health concept, which emphasizes that antimicrobial resistance circulates across clinical, agricultural, and environmental systems through wastewater networks, horizontal gene transfer, and other environmental pathways.

**Table 2:** Common reservoirs of the 15 ARG resistance classes reported in the literature. These associations reflect general patterns described in prior wastewater, clinical, and environmental surveillance studies and are presented as interpretive context rather than as a quantitative attribution of provenance within the sampled cities.

| ARG Class | Common reservoirs reported in literature | Reference |
| --- | --- | --- |
| Aminoglycoside (AMG) | Livestock wastewater, hospital wastewater | [79, 80] |
| β-lactam (BL) | Hospital wastewater, clinical isolates | [80, 81] |
| Tetracycline (TET) | Livestock, manure, agricultural soils, wastewater | [79, 80] |
| Sulfonamide (SUL) | Wastewater, municipal sewage, rivers | [82, 83] |
| Macrolide–lincosamide–streptogramin (MLS) | Livestock, humans, wastewater | [79, 80] |
| Trimethoprim (TMP) | Wastewater, human gut, clinical settings | [80, 84] |
| Fluoroquinolone (FQ) | Hospital wastewater, municipal wastewater | [84] |
| Phenicol (PHEN) | Livestock, aquaculture, wastewater | [84, 80] |
| Glycopeptide (GLY) | Hospitals (*Enterococcus*), historical livestock use (avoparcin) | [85] |
| Fosfomycin (FOS) | Hospitals, urinary tract isolates, wastewater | [80, 84] |
| Colistin (COL) | Livestock (historical veterinary use), wastewater, environment | [86] |
| Nitroimidazole (NI) | Human gut, anaerobic bacteria | [84, 80] |
| Oxazolidinone (OXA) | Livestock (*Enterococcus*), hospital-associated clades | [87, 84] |
| Rifamycin (RIF) | Environmental/soil bacteria (intrinsic), clinical settings | [84] |
| Multidrug/efflux pump-associated (MDR-EP) | Ubiquitous (intrinsic, environmental and clinical bacteria) | [80] |

### 4.2. Geographic Structuring of the Urban Resistome

Principal Coordinates Analysis (PCoA) based on Bray–Curtis dissimilarity (Figure 4) revealed clear geographic clustering of wastewater resistomes, while PERMANOVA confirmed significant differences in resistome composition among cities (F = 68.59, R² = 0.54, p = 0.001). These findings indicate that ARG communities differed beyond random variation and suggest that local environmental conditions, anthropogenic activities, and wastewater management practices collectively contribute to regional resistome composition [88, 89].

City identity accounted for approximately 54% of the variation in resistome composition, leaving 46% attributable to within-city variability. This result suggests that models capable of transferring across cities are likely capturing environmental relationships that extend beyond city identity alone. The observed clustering is also consistent with the possibility that differences in microbial community structure and gene exchange dynamics contribute to the spatial heterogeneity of ARG distributions.

Several ARG classes investigated in this study are frequently associated with mobile genetic elements (MGEs) that facilitate horizontal gene transfer among bacterial populations [90]. For example, sulfonamide resistance genes (*sul1*, *sul2*) are commonly linked to class 1 integrons [91], while *β*-lactamase genes (*bla*), colistin resistance genes (*mcr*), and tetracycline resistance genes (*tetA*, *tetM*, *tetO*) are frequently carried on conjugative plasmids and transposons [90]. Consequently, regional variation in MGE prevalence and transmission dynamics may partly explain the geographic clustering identified by PERMANOVA. Although MGEs were not directly quantified in the present study, future work integrating metagenomic characterization of plasmids, integrons, and transposons with explainable machine learning could provide mechanistic insight into the interactions among environmental conditions, horizontal gene transfer, and ARG dissemination.

This hypothesis also generates a prediction that can be explored using the present dataset. If mobilization enhances geographic transferability, ARG classes predominantly associated with promiscuous, well-conserved mobile genetic elements should exhibit greater cross-city predictive performance than classes whose distributions are primarily governed by local selection. The per-class classification results presented in Section 4.4 are consistent with this expectation. Sulfonamide resistance (ROC–AUC = 0.976), largely represented by *sul1* and *sul2* associated with class 1 integrons, and *β*-lactam resistance (ROC–AUC = 0.981), frequently associated with conjugative plasmids, showed the highest transferability across cities. In contrast, glycopeptide resistance exhibited the lowest predictive performance (ROC–AUC = 0.762), consistent with its predominantly chromosomal distribution and comparatively limited mobilization potential. Although these observations are consistent with the proposed hypothesis, they do not establish a causal relationship. Direct quantification of integrons, plasmids, and transposons within the same wastewater samples will be necessary to determine whether mobilization potential predicts geographic transferability, as discussed in Section 5.

### 4.3. Model Performance Comparison

Four supervised machine learning algorithms, LR, RF, XGBoost, and CatBoost, were systematically evaluated to identify the most effective framework for predicting ARG occurrence across geographically distinct urban wastewater systems. Model performance was assessed using LOGO crossvalidation to evaluate generalizability to previously unseen cities.

As summarized in Table 3 and Figure 5, CatBoost achieved the best overall predictive performance, yielding the highest median ROC–AUC (0.929), mean ROC–AUC (0.877), and median F1-score (0.750). Logistic Regression and Random Forest also demonstrated competitive performance, with Random Forest achieving a median ROC–AUC of 0.917 and a mean ROC–AUC of 0.866.

**Table 3:** Performance comparison of machine learning models for ARG detectionoccurrence prediction under LOGO cross-validation (n = 5 folds).

| Metric | CatBoost | Logistic Regression | Random Forest | XGBoost |
| --- | --- | --- | --- | --- |
| ROC-AUC Med. | 0.929 | 0.900 | 0.917 | 0.889 |
| ROC-AUC Mean | 0.877 | 0.859 | 0.866 | 0.827 |
| ROC-AUC Std. | 0.163 | 0.126 | 0.161 | 0.204 |
| F1 Mean | 0.660 | 0.600 | 0.669 | 0.659 |
| F1 Median | 0.750 | 0.700 | 0.741 | 0.741 |

**Figure 5:**
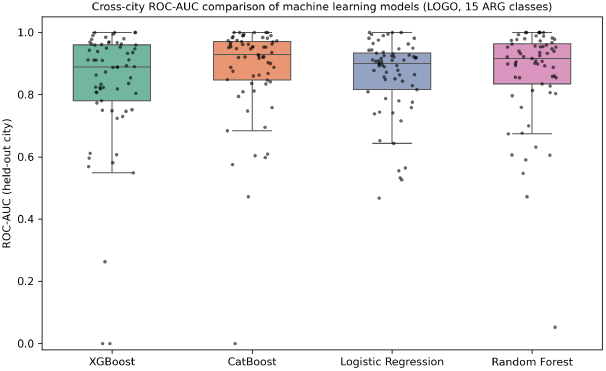
Cross-city ROC-AUC comparison of machine learning models.

The performance advantage of CatBoost over the linear baseline was modest (median ROC-AUC: 0.929 vs. 0.900; mean ROC-AUC: 0.877 vs. 0.859), indicating that much of the predictive signal was captured by a relatively simple linear model, while gradient boosting provided an incremental improvement by modeling nonlinear relationships and feature interactions.

Although CatBoost achieved the highest overall predictive accuracy, its performance varied across the LOGO folds (ROC-AUC SD = 0.163), indicating that predictive performance varied on the held-out city. This geographic variability likely reflects differences in local environmental conditions, resistome composition, and sampling characteristics, and is examined in greater detail in Section 4.4.

Figure 5 further illustrates the distribution of ROC–AUC values across ARG classes and study cities. CatBoost maintained consistently high predictive performance across most city–ARG combinations, whereas the remaining models exhibited greater variability among validation folds, indicating reduced stability when applied to previously unseen geographic locations.

The superior performance of CatBoost is likely attributable to its ability to capture nonlinear relationships and feature interactions among environmental and omics-derived predictors. Its strong performance under LOGO cross-validation further indicates that these relationships generalize beyond the cities included in model training, supporting the suitability of the proposed framework for geographically transferable environmental surveillance.

Table 4 summarizes the effect of validation strategy and feature selection on CatBoost performance. Changing from random 5-fold cross-validation to geographically independent LOGO validation reduced ROC–AUC by 0.052, whereas removing geographic coordinates reduced ROC–AUC by only 0.003 under 5-fold cross-validation. Removing the metagenomic feature block reduced ROC–AUC by 0.384 relative to the full-feature LOGO model. Together, these comparisons indicate that geographic transferability imposed a larger performance penalty than either explicit geographic coordinates or the omission of metagenomic features.

**Table 4:** Effect of validation strategy and feature selection on CatBoost performance.

| Design | Configuration | ROC-AUC | $\Delta$ | Interpretation |
| --- | --- | --- | --- | --- |
| Reference | Full, 5-fold | $0.929 \pm 0.061$ | – | Baseline |
| Validation | Full, LOGO | $0.877 \pm 0.163$ | –0.052 | Transfer penalty |
| Coordinates | Nolat/lon, 5-fold | $0.926 \pm 0.062$ | –0.003 | Coordinate effect |
| Validation+<br>Coordi-<br>nates | Nolat/lon,<br>LOGO | $0.870 \pm 0.158$ | –0.059 | Transfer without coordinates |
| Omics | Abiotic<br>only(LOGO) | $0.493 \pm 0.159$ | –0.384<sup>a</sup> | Sequencing contribution |

<sup>a</sup> Difference relative to the full-feature LOGO model (0.877).

### 4.4. Per-Class ARG Prediction Performance

Although CatBoost achieved the highest overall predictive performance, classification accuracy varied substantially among individual ARG classes (Table 5 and Figure 6). The best-performing classes were *β*-lactam (BL; median ROC–AUC = 0.981, SD = 0.015), sulfonamide (SUL; 0.976, SD = 0.018), aminoglycoside (AMG; 0.975, SD = 0.071), and phenicol (PHEN; 0.954, SD = 0.029). These classes also achieved the highest median F1-scores (0.815–0.875), indicating consistently reliable classification across geographically independent validation folds.

**Table 5:**
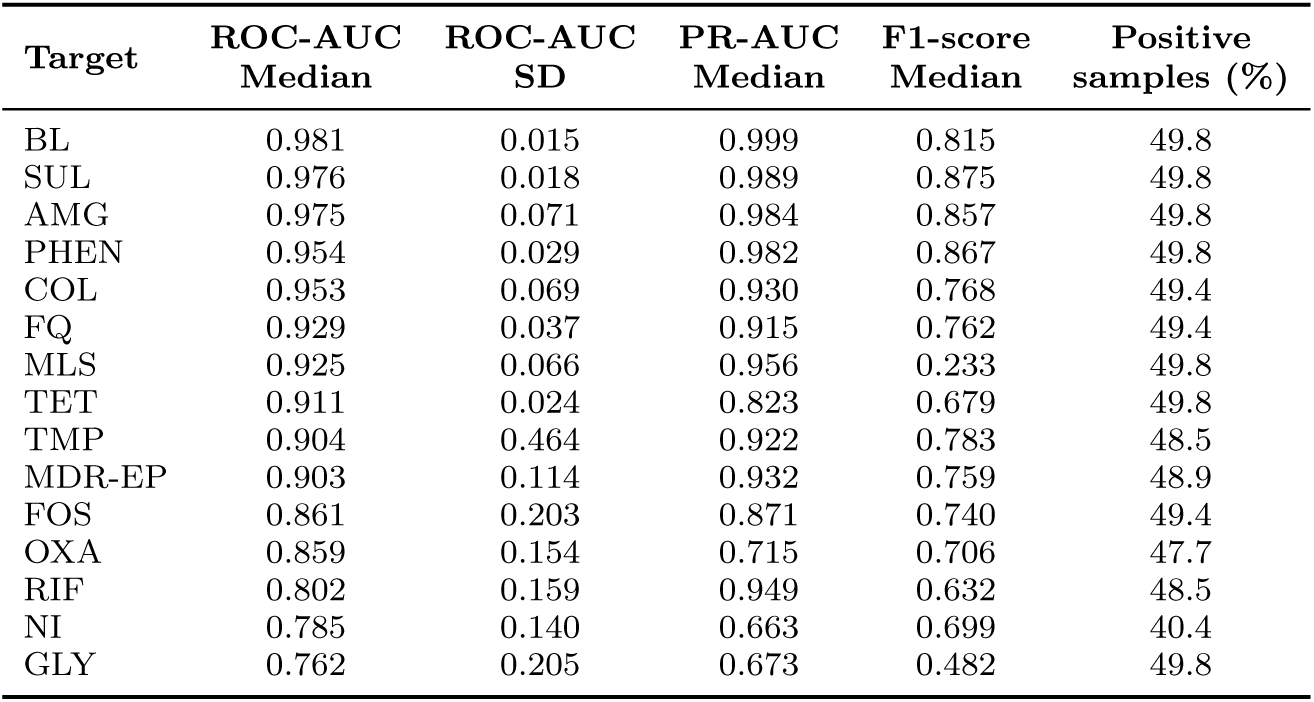
Predictive performance of the CatBoost model for each target resistance class under leave-one-group-out (LOGO) cross-validation. ROC-AUC and PR-AUC are median values across the five LOGO folds; SD is the standard deviation of ROC-AUC across the held-out cities; positive samples is the proportion assigned to the positive class after median-threshold binarization.

| Target | ROC-AUC<br>Median | ROC-AUC<br>SD | PR-AUC<br>Median | F1-score<br>Median | Positive<br>samples (%) |
| --- | --- | --- | --- | --- | --- |
| BL | 0.981 | 0.015 | 0.999 | 0.815 | 49.8 |
| SUL | 0.976 | 0.018 | 0.989 | 0.875 | 49.8 |
| AMG | 0.975 | 0.071 | 0.984 | 0.857 | 49.8 |
| PHEN | 0.954 | 0.029 | 0.982 | 0.867 | 49.8 |
| COL | 0.953 | 0.069 | 0.930 | 0.768 | 49.4 |
| FQ | 0.929 | 0.037 | 0.915 | 0.762 | 49.4 |
| MLS | 0.925 | 0.066 | 0.956 | 0.233 | 49.8 |
| TET | 0.911 | 0.024 | 0.823 | 0.679 | 49.8 |
| TMP | 0.904 | 0.464 | 0.922 | 0.783 | 48.5 |
| MDR-EP | 0.903 | 0.114 | 0.932 | 0.759 | 48.9 |
| FOS | 0.861 | 0.203 | 0.871 | 0.740 | 49.4 |
| OXA | 0.859 | 0.154 | 0.715 | 0.706 | 47.7 |
| RIF | 0.802 | 0.159 | 0.949 | 0.632 | 48.5 |
| NI | 0.785 | 0.140 | 0.663 | 0.699 | 40.4 |
| GLY | 0.762 | 0.205 | 0.673 | 0.482 | 49.8 |

**Figure 6:**
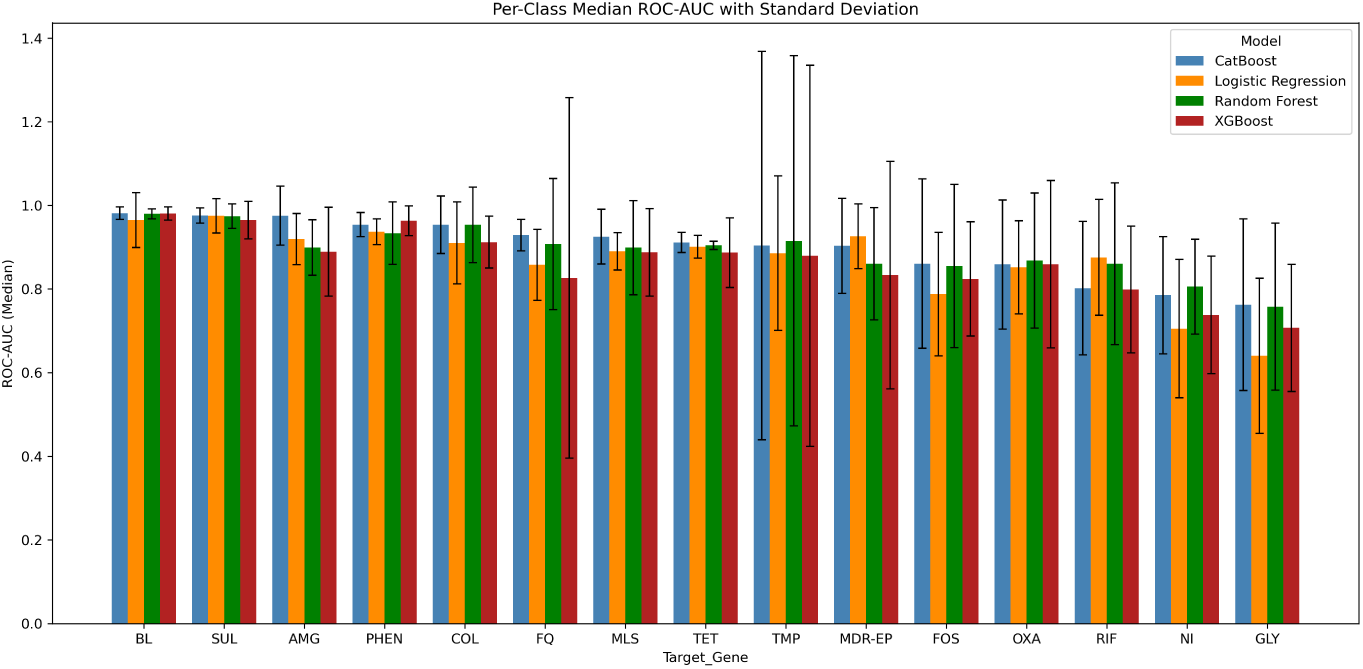
Per-class ROC-AUC of the four machine learning models under leave-one-group-out (LOGO) cross-validation. Error bars represent the standard deviation across the five held-out cities.

Intermediate predictive performance was observed for colistin (COL), fluoroquinolone (FQ), macrolide–lincosamide–streptogramin (MLS), tetracycline (TET), trimethoprim (TMP), and multidrug/efflux pump-associated genes (MDR-EP), with median ROC–AUC values ranging from 0.903 to 0.953. However, predictive stability varied considerably among these classes. COL, FQ, and TET exhibited relatively low variability (SD *≤* 0.069), whereas TMP showed the largest variability among all resistance classes (SD = 0.464), despite a median ROC–AUC of 0.904. The large discrepancy between the median ROC–AUC (0.904) and the corresponding mean ROC–AUC (0.695) indicates that TMP predictions were highly inconsistent across the five heldout cities and should therefore not be considered consistently transferable based on median performance alone.

Lower predictive performance was observed for fosfomycin (FOS), oxazolidinone (OXA), rifamycin (RIF), nitroimidazole (NI), and glycopeptide (GLY), with median ROC–AUC values ranging from 0.762 to 0.861 and relatively high variability among validation folds (SD = 0.140–0.205). Several of these classes were also among the less prevalent targets (Table 5), suggesting that limited positive samples may have contributed to reduced geographic transferability through increased statistical uncertainty.

MLS resistance exhibited a different limitation. Although MLS achieved a relatively high median ROC–AUC (0.925), it produced the lowest median F1-score (0.233), indicating that the model ranked positive samples accurately while the fixed classification threshold transferred poorly across cities. Consequently, MLS predictions may be more appropriate for relative risk ranking than binary classification.

Overall, class-specific transferability varied considerably, with a 0.219 difference in median ROC–AUC between *β*-lactam (0.981) and glycopeptide (0.762). These results demonstrate that predictive performance should be evaluated separately for each ARG class rather than relying solely on an overall model summary. The observed performance gradient is consistent with the hypothesis discussed in Section 4.2 that ARG classes influenced by broadly shared environmental processes exhibit greater geographic transferability, although this interpretation remains a testable hypothesis because mobile genetic elements were not directly quantified.

### 4.5. Statistical Comparison of Model Performance

To assess whether the performance improvements of CatBoost over baseline algorithms were statistically significant, the Wilcoxon signed-rank test was applied to ROC-AUC scores across LOGO folds. This non-parametric test was selected for its robustness to non-normal distributions and its suitability for paired comparisons across multiple evaluation folds. Pairwise Wilcoxon signed-rank tests confirmed that CatBoost significantly outperformed XGBoost (W=272.5, p<0.001), and Logistic Regression (W=538.0, p=0.002). These results demonstrate that the superior predictive performance of CatBoost is statistically significant and unlikely to be attributable to random variation or specific geographic outliers.

### 4.6. SHAP Analysis of Abiotic Determinants of ARG Occurrence

To identify the abiotic predictors contributing to ARG occurrence across the five study cities, SHAP analysis was performed for the overall dataset and individually for each city (Figure 7). Across all cities, latitude and longitude consistently exhibited the highest SHAP values, indicating that broad geographic gradients contributed substantially to model predictions. Surface pressure (PS), wind speed (WS2M), evapotranspiration (EVPTRNS), and other atmospheric and hydrological variables were also among the most influential predictors, indicating that the model relied on multiple abiotic factors beyond geographic location alone.

**Figure 7:**
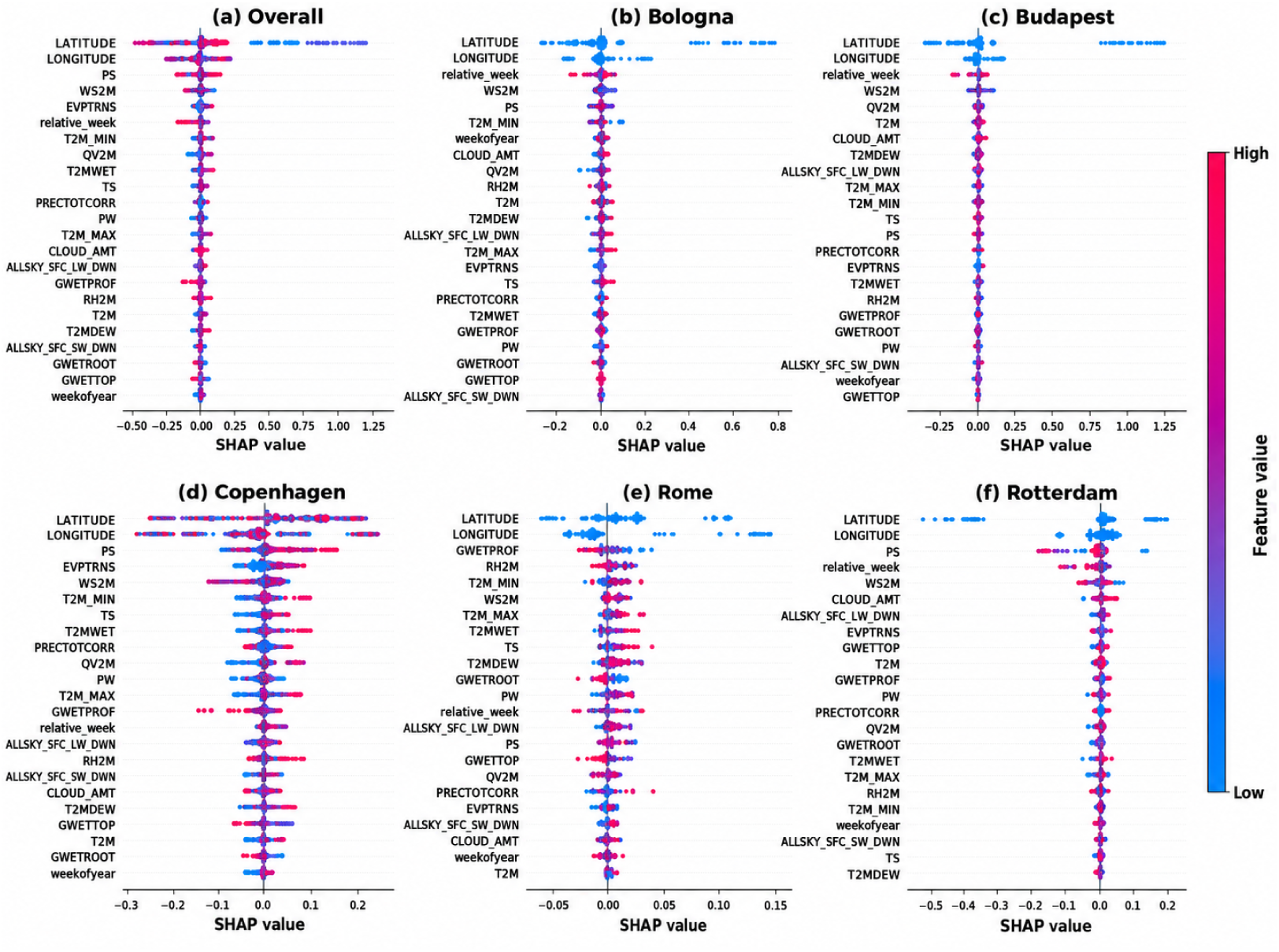
SHAP-based interpretation of the CatBoost model. Beeswarm plots of the SHAP contributions of the 23 abiotic predictors to ARG prediction for (a) the overall dataset, (b) Bologna, (c) Budapest, (d) Copenhagen, (e) Rome and (f) Rotterdam. Positive and negative SHAP values indicate increased and decreased contributions to the predicted probability of ARG occurrence; point colour gives the relative value of each predictor.

The SHAP rankings were consistent with the environmental processes discussed in Section 2.3. While geographic coordinates captured broad spatial gradients, atmospheric and hydrological variables were consistently identified as dominant contributors, supporting the relevance of transport- and persistence-related processes. Temperature-related variables exhibited comparatively lower SHAP importance, likely because part of their predictive information was shared with latitude and longitude across the geographic gradient represented by the five cities. Their lower SHAP values therefore should not be interpreted as evidence that temperature plays only a limited role in ARG occurrence.

City-specific SHAP patterns further demonstrated environmental heterogeneity. Copenhagen and Rotterdam exhibited stronger contributions from atmospheric and hydrological variables, whereas Rome and Bologna showed relatively greater contributions from temperature- and moisture-related predictors. Variance analysis found no significant relationship between standardized predictor variance and city-level SHAP importance (Spearman *ρ* = *−*0.32, *p* = 0.084), although the statistical power was limited by the inclusion of only five cities (Table S5 and Figure S2). Notably, latitude and longitude exhibited negligible within-city variance while remaining the most influential predictors overall, indicating that their importance primarily reflects differences among cities rather than within-city variability. These differences therefore likely reflect environmental variation among cities rather than distinct underlying mechanisms.

The observed SHAP patterns are consistent with previous studies showing that large-scale geographic gradients reflect differences in climate, land use, population density, and wastewater management across European cities [88]. Similarly, the importance of surface pressure, wind speed, and evapotranspiration agrees with their reported influence on microbial transport, deposition, environmental persistence, and hydrological processes [65, 63, 64]. Although SHAP does not establish causality, the agreement between model interpretation and established environmental processes supports the biological plausibility of the learned relationships.

### 4.7. Independent Predictive Contribution of Environmental Variables

To evaluate the predictive contribution of environmental variables, all ML models were retrained using only abiotic predictors, excluding all omics-derived features. As summarized in Table 6, predictive performance decreased for all models relative to the integrated framework; however, CatBoost continued to achieve the highest performance using environmental predictors alone.

**Table 6:**
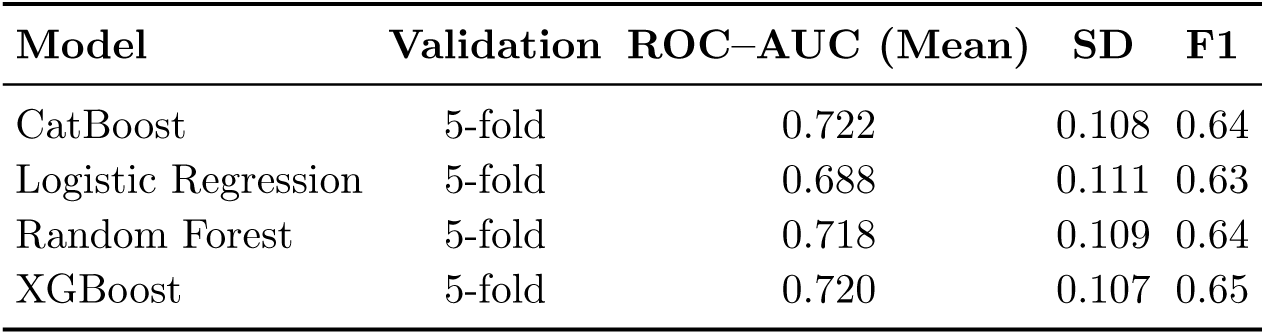
Model performance using the 23 abiotic predictors only (no omics-derived features) under random stratified 5-fold cross-validation. ROC–AUC is averaged over 75 (5-fold) ARG class *×* fold evaluations, F1 over all 75; SD is the standard deviation of ROC–AUC across evaluations.

| Model | Validation | ROC-AUC (Mean) | SD | F1 |
| --- | --- | --- | --- | --- |
| CatBoost | 5-fold | 0.722 | 0.108 | 0.64 |
| Logistic Regression | 5-fold | 0.688 | 0.111 | 0.63 |
| Random Forest | 5-fold | 0.718 | 0.109 | 0.64 |
| XGBoost | 5-fold | 0.720 | 0.107 | 0.65 |

Although predictive performance decreased after excluding omics-derived features, all four models retained substantial predictive capability, indicating that abiotic variables contain meaningful information associated with ARG occurrence. CatBoost achieved the highest ROC–AUC (0.722), suggesting that nonlinear relationships among environmental predictors provide considerable discriminatory power even without metagenomic information. Expressed relative to the integrated framework (mean ROC–AUC = 0.887), the abiotic-only CatBoost model recovered 78% of the full model’s predictive signal, using only freely available environmental reanalysis data spanning meteorological, hydrological, radiative, geospatial, and temporal domains; this is consistent with prior work demonstrating that environmental and climatic variables serve as effective proxies for ARG occurrence and abundance in the absence of direct resistome profiling [52, 62]. The abiotic-only results reported here can be placed alongside the beach-water ARG study of Zahra et al. [62], which modelled qPCR-derived ARG abundance from six hydrometeorological predictors (water temperature, tide, salinity, precipitation, wind speed and air pressure) with CatBoost under a random train–test split. Compared with the integrated including full features (mean ROC–AUC = 0.877), the reduction in predictive performance confirms that metagenomic features provide complementary biological information beyond that captured by environmental variables. Nevertheless, the relatively strong performance obtained using only abiotic predictors is consistent with the SHAP analysis of the integrated model, which identified atmospheric, hydrological, and geospatial variables as major contributors to model predictions.

These findings demonstrate that environmental predictors alone provide substantial predictive information for ARG occurrence while also highlighting the complementary value of integrating environmental and metagenomic information. From an environmental monitoring perspective, these results suggest that abiotic variables can support preliminary ARG risk assessment where routine metagenomic sequencing is unavailable or economically impractical, whereas the highest predictive performance is achieved by integrating environmental and metagenomic information.

### 4.8. Geographic Feature Sensitivity Analysis

The SHAP analysis identified latitude and longitude as two of the most influential predictors of ARG occurrence (Figure 7). Although geographic coordinates capture broad spatial variability, their high SHAP importance raises the possibility that predictive performance may partly reflect spatial autocorrelation rather than transferable environmental relationships. To evaluate the contribution of geographic information, latitude and longitude were removed from the feature set, and CatBoost was re-evaluated using both 5-fold cross-validation and LOGO cross-validation.

As summarized in Table 7, removing geographic coordinates resulted in only modest reductions in mean ROC–AUC under both validation regimes. Under 5-fold cross-validation, mean ROC–AUC decreased from 0.929 *±* 0.061 (full features) to 0.926 *±* 0.062 (without coordinates), whereas under LOGO cross-validation it decreased from 0.877 *±* 0.163 to 0.870 *±* 0.158. The F1score showed a similarly small decrease under 5-fold cross-validation (0.847 *±* 0.078 to 0.842 *±* 0.078), but a modest increase under LOGO cross-validation (0.660 *±* 0.315 to 0.655 *±* 0.318). Overall, these results indicate that removing latitude and longitude had only a modest effect on predictive performance, suggesting that the remaining abiotic and omics-derived predictors retained most of the information required for ARG prediction.

**Table 7:** Geographic feature sensitivity analysis of CatBoost performance.

| Feature Set | Validation | ROC–AUC | F1 Score | $\Delta$ ROC–AUC vs Full |
| --- | --- | --- | --- | --- |
| Full Features | 5-Fold CV | $0.929 \pm 0.061$ | $0.847 \pm 0.078$ | – |
| Full Features | LOGO | $0.877 \pm 0.163$ | $0.660 \pm 0.315$ | –0.053 |
| Without Coordinates | 5-Fold CV | $0.926 \pm 0.062$ | $0.842 \pm 0.078$ | –0.003 |
| Without Coordinates | LOGO | $0.870 \pm 0.158$ | $0.655 \pm 0.318$ | –0.007 |

Notably, the difference in predictive performance between the two validation strategies was considerably larger than the effect of removing geographic coordinates. Consequently, differences among urban environments, rather than dependence on geographic coordinates alone, appear to represent the principal challenge for robust model generalization. The modest reduction in predictive performance after removing latitude and longitude therefore further supports the robustness of the integrated modeling framework and suggests that the model learned relationships between environmental conditions and ARG occurrence that extend beyond explicit geographic coordinates. Comprehensive performance comparisons for all evaluated classifiers under both validation strategies and feature configurations are provided in Table S6, while the corresponding ROC–AUC and F1-score comparisons are shown in Figure S3.

### 4.9. Geospatial Prediction of High-Risk ARG Occurrence Across Urban Wastewater Systems

To demonstrate the practical applicability of the proposed framework, the proportion of predicted high-risk ARG occurrences was calculated for each city by averaging the binary CatBoost predictions generated under LOGO cross-validation across all wastewater samples and the 15 ARG resistance classes. These city-level proportions were subsequently visualized geographically (Figure 8a).

**Figure 8:**
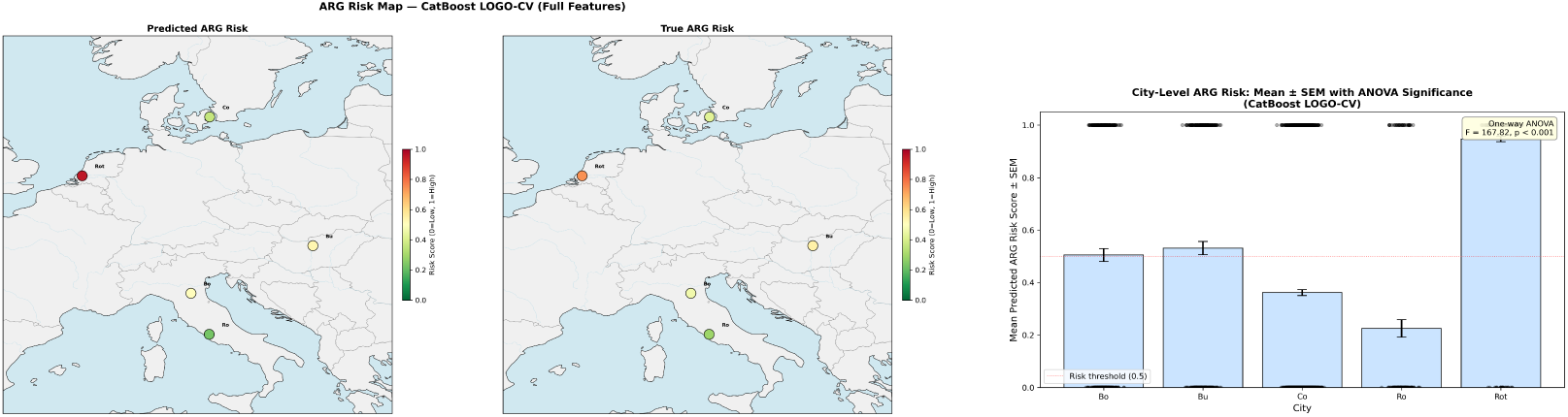
Geospatial ARG risk assessment across the five study cities. (a) Predicted (left) and observed (right) composite ARG risk scores derived from CatBoost LOGO cross-validation, mapped across the five European study cities. Each point is colored according to the proportion of predicted high-risk ARG occurrences aggregated across all 15 resistance classes. (b) City-level predicted ARG risk (mean *±* SEM) with one-way ANOVA significance annotation.

The spatial distribution of predicted high-risk ARG occurrence closely mirrored the observed geographic patterns across the five study cities. Rotterdam exhibited the highest proportion of predicted high-risk ARG occurrences, whereas Rome showed the lowest. Bologna and Budapest displayed comparable intermediate levels, while Copenhagen exhibited a lower proportion than both cities. This agreement between predicted and observed spatial patterns suggests that the CatBoost framework successfully captured the major geographic differences in ARG occurrence, even when an entire city was excluded during model training.

One-way ANOVA (Table S7) identified significant differences in the proportion of predicted high-risk ARG occurrences among cities (*F* = 167.82, *p <* 0.001; Figure 8b). Tukey’s HSD post-hoc analysis showed that nine of the ten pairwise city comparisons were statistically significant (*p <* 0.05), with Bologna and Budapest representing the only non-significant comparison (*p* = 0.927). Rotterdam exhibited the highest mean predicted proportion (0.947 *±* 0.010, standard error of the mean (SEM)), whereas Rome exhibited the lowest (0.225 *±* 0.033, SEM). Complete summary statistics are provided in Tables S7 and S8.

From an environmental surveillance perspective, the geographic visualizations illustrate how held-out LOGO predictions can be summarized to prioritize locations for further investigation. However, these maps should not be interpreted as validated risk maps or operational decision-support products, because the framework has not undergone prospective field validation, probability calibration, or evaluation beyond the five study cities. ARG-class-specific prediction maps are presented in Figure S4 and Figure S5.

### 4.10. One Health Implications

The proposed framework has important implications within the One Health concept, which recognizes the interconnected nature of human, animal, and environmental health in addressing antimicrobial resistance (AMR) [88]. By integrating metagenomic observations with abiotic environmental predictors, the framework shows that environmental conditions contain substantial information for predicting ARG occurrence across geographically diverse urban wastewater systems. This highlights the value of combining biological and environmental information to strengthen environmental AMR surveillance.

Several observations support this integrated One Health perspective. CatBoost maintained strong predictive performance under geographically independent LOGO validation, indicating that the learned relationships generalize beyond individual cities. SHAP analysis consistently identified atmospheric, hydrological, temporal, and geospatial variables as important contributors to model predictions, while the environmental-only models retained considerable predictive capability. Furthermore, removing latitude and longitude resulted in only modest reductions in predictive performance, indicating that the model relied on multiple environmental predictors rather than explicit geographic coordinates alone.

From a practical perspective, all abiotic predictors used in this study were obtained from the freely accessible NASA POWER platform, which provides global gridded meteorological and environmental reanalysis data through an open API without requiring local monitoring infrastructure. Consequently, the framework can be applied in regions where dense national meteorological observation networks are unavailable, improving its scalability and accessibility. When combined with metagenomic observations, these globally available environmental data provide a practical approach for supporting ARG surveillance, particularly in regions where routine metagenomic sequencing remains financially or technically challenging. Continuous environmental predictions can therefore support preliminary ARG risk assessment and help prioritize locations and time periods for targeted metagenomic sampling.

The One Health relevance of the proposed framework is further strengthened by the ecological context presented in Section 4.1. As discussed there, the ARG classes detected in this study have been widely reported across interconnected human, animal, and environmental reservoirs, providing ecological context for the wastewater resistome. Within this context, the ability to estimate ARG occurrence from abiotic environmental conditions further supports the use of wastewater as an integrated One Health surveillance matrix. However, the model predicts ARG occurrence in wastewater as an indicator of environmental exposure rather than antimicrobial resistance transmission, resistance phenotypes, or clinical disease burden. It should therefore be regarded as a decision-support tool for environmental surveillance and sampling prioritization that complements, rather than replaces, conventional metagenomic surveillance. By integrating environmental and metagenomic information within a geographically transferable framework, the proposed approach contributes to more scalable environmental AMR surveillance and supports the implementation of One Health monitoring strategies, including those advocated by the WHO Global Action Plan on Antimicrobial Resistance [6].

## 5. Surveillance Deployment Readiness

The framework proposed here is intended to guide environmental monitoring efforts, which requires explicitly defining the conditions under which its predictions are appropriate for practical use. NASA Technology Readiness Levels (TRLs) were developed to assess aerospace hardware and are not directly applicable to predictive machine learning models, which are instead characterized by attributes such as data latency, cost per decision-grade prediction, geographic transferability, interpretability, and applicability domain. Accordingly, Table 8 introduces a Surveillance Deployment Readiness (SDR) framework that evaluates the proposed approach across these five dimensions and summarizes its readiness for environmental surveillance applications.

**Table 8:**
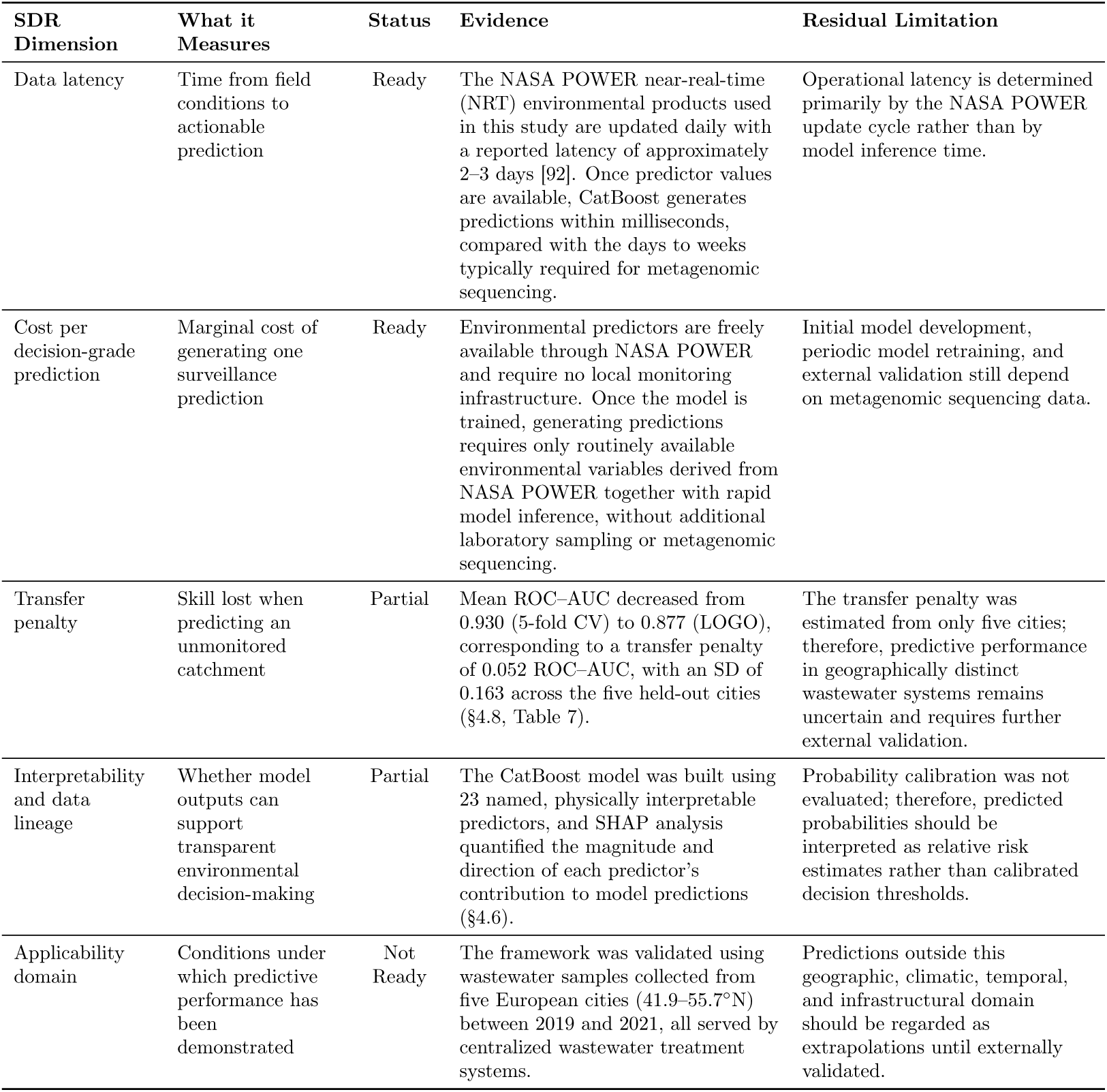
Surveillance Deployment Readiness (SDR) assessment of the proposed framework.

The proposed framework is intended to complement rather than replace conventional metagenomic surveillance. Metagenomic sequencing remains essential for direct ARG detection, identification of novel resistance determinants, and comprehensive characterization of wastewater resistomes. In contrast, the abiotic-only framework developed in this study provides rapid environmental risk screening using routinely available environmental variables without requiring additional sequencing after model deployment. Together, these complementary approaches enable continuous environmental screening while reserving metagenomic sequencing for targeted confirmation and detailed resistome characterization.

A comparison of conventional metagenomic surveillance, the integrated prediction framework, and the proposed abiotic-only framework is provided in Table S9.

Table 9 summarizes the principal conditions under which the framework’s predictions should be interpreted cautiously or not relied upon. For each failure mode, the table identifies the underlying mechanism, the observable signature where one is available, the recommended mitigation strategy, and the residual risk that remains after mitigation.

**Table 9:** Prediction failure modes, observable signatures, mitigation strategies, and residual risks of the proposed framework.

| Failure mode | Mechanism | Observable signature | Mitigation | Residual risk |
| --- | --- | --- | --- | --- |
| Rare-class label uncertainty | Assembly-based metagenomic workflows may fail to recover some low-abundance ARGs, introducing label uncertainty that is likely to be greatest for rare resistance classes (§3.2). | Lower ROC-AUC and greater fold-to-fold variability were observed for several of the least prevalent classes (e.g., GLY, NI, and RIF). | Report class prevalence for each city; consider read-mapping approaches for rare determinants; interpret predictions cautiously for low-prevalence classes; and abstain from binary classification when prevalence is insufficient for reliable validation. | Prediction uncertainty is greatest for rare ARG classes, which may include clinically important or emerging resistance determinants. |
| Decision-threshold miscalibration | Classification thresholds optimized using the training cities may not remain optimal in wastewater systems with different ARG prevalence or class distributions. | High ROC-AUC accompanied by substantially lower F1-score (e.g., MLS), indicating reliable ranking but unstable binary classification. | Where possible, recalibrate decision thresholds using a representative labelled subset from the target site; otherwise, interpret predictions primarily as relative risk rankings rather than binary classifications. | Threshold recalibration requires additional labelled data and therefore cannot eliminate the need for periodic metagenomic sequencing. |
| Out-of-domain extrapolation | The framework may be applied to wastewater systems outside the climatic, geographic, temporal, or infrastructural conditions represented by the five training cities. | No direct performance signature exists without external labelled data; the model may return apparently reliable probability estimates even for samples outside the validated domain. | Define an applicability domain using the environmental predictor space; flag samples outside this domain; and abstain from prediction until external validation is available. | Performance in substantially different wastewater systems remains unknown, making this the most important limitation for operational deployment. |
| Spatial resolution limitation | NASA POWER predictors represent gridded environmental conditions whose spatial resolution may exceed that of individual wastewater treatment plants or catchments (§3.1.1). | Nearby wastewater treatment plants represented by the same or similar grid cells may receive nearly identical environmental predictor values. | Report the spatial resolution of the environmental predictors explicitly; avoid plant-level interpretation where multiple facilities share similar gridded inputs; and use local meteorological or catchment measurements when finer spatial discrimination is required. | Plant-level environmental differences may not be fully resolved when multiple facilities are represented by similar gridded environmental data. |
| Temporal drift | Changes in antimicrobial use, wastewater treatment practices, infrastructure, climate, or microbial ecology may alter the relationship between environmental predictors and ARG occurrence after the 2019–2021 study period. | Model degradation cannot be detected directly from environmental predictors alone; shifts in predictor or prediction distributions may indicate potential drift. | Periodically revalidate the framework against a representative sequenced subsample (e.g., annually), together with monitoring of predictor and prediction distributions for evidence of dataset drift. | The appropriate revalidation interval cannot be estimated from the three-year dataset; the proposed annual schedule represents a practical operational recommendation rather than an empirically derived interval. |
| Annotation database limitation | ARG labels were derived using CARD-based annotation; consequently, divergent or previously uncharacterized resistance determinants not represented in the reference database cannot be included reliably in the training labels (§3.2). | Novel resistance determinants are absent from both the training labels and model predictions, with no explicit warning generated by the framework. 38 | State explicitly that predictions are limited to ARG classes represented in the training data and CARD annotation workflow; periodically update annotations using newer CARD releases and retrain the model when appropriate. | The framework cannot identify emerging ARGs until they are incorporated into updated reference databases and future training datasets. |

Taken together, Tables 8 and 9 define the operational scope of the proposed framework. The framework is best suited for routine environmental risk screening in wastewater systems that resemble the climatic, geographic, temporal, and infrastructural conditions represented by the five study cities. However, predictions should be interpreted cautiously for rare ARG classes, threshold-sensitive classifications, samples outside the validated applicability domain, and monitoring periods affected by temporal drift.

## 6. Conclusion

This study investigated how much of the predictive signal for ARG occurrence in urban wastewater remained after excluding an entire city from model training and how much of that predictive capability could be retained without metagenomic sequencing. Changing the validation strategy from 5-fold cross-validation to geographically independent LOGO validation reduced ROC–AUC by 0.052, whereas removing geographic coordinates reduced ROC–AUC by only 0.003 under 5-fold cross-validation, indicating that the effect of geographic transferability exceeded the contribution of explicit geographic coordinates. Removing all metagenomic features reduced the CatBoost ROC–AUC from 0.877 to 0.722, while the abiotic-only model still achieved strong predictive performance using freely available environmental reanalysis data.

Model performance also varied among ARG classes, with ROC–AUC values ranging from 0.762 for glycopeptide resistance genes to 0.981 for *β*-lactam resistance genes. This variability suggests that geographically transferable prediction may depend on the underlying environmental processes associated with different resistance classes. SHAP analysis indicated that the highest-performing classes relied more strongly on atmospheric, hydrological, and temporal predictors, whereas lower-performing classes showed greater dependence on geographic coordinates. Together, these observations suggest that geographically transferable prediction is more likely when model decisions are driven by broadly shared environmental processes rather than explicit geographic location.

The results indicate that freely available environmental reanalysis data can support rapid, low-cost prediction of ARG occurrence risk and help prioritize wastewater catchments for targeted metagenomic sequencing. This capability was strongest for the highest-performing resistance classes and for wastewater systems resembling the five cities included in the present study. However, the framework is not a substitute for metagenomic sequencing, as it cannot directly detect ARGs, identify previously uncharacterized resistance determinants, or predict resistance beyond the reference database used for model training. Furthermore, performance outside the validated geographic and environmental domain remains uncertain, emphasizing the importance of external validation before operational deployment.

The conclusions of this study are necessarily conditional on the environmental and geographic domain represented by the available data. The results suggest that geographically transferable prediction is achievable when the relationship between environmental conditions and ARG occurrence is captured by environmental variables that generalize across locations. Conversely, predictive performance is expected to decline where these relationships differ substantially from those represented in the training data. Future prospective validation across diverse wastewater systems will therefore be essential to establish the operational applicability of the proposed framework.

## CRediT authorship contribution statement

**Somayeh Falahati Khanaman:** Conceptualization, Methodology, Formal analysis, Visualization, Writing – original draft, Writing – review & editing; **Etienne Gnimpieba:** Conceptualization, Methodology, Data curation, Formal analysis, Writing – review & editing, Supervision; **Bichar Dip Shrestha Gurung:** Data curation; **Shiva Aryal:** Methodology; **Mengistu Geza Nisrani:** Supervision; **Venkataramana Gadhamshetty:** Conceptualization, Writing – review & editing, Supervision, Project administration, Funding acquisition.

## Declaration of generative AI and AI-assisted technologies in the writing process

During the preparation of this work, the author(s) used [Overleaf AI, Claude Code and ChatGPT TOOL/SERVICE] for [reviewing and refining]. The author(s) reviewed and edited the output as needed and took full responsibility for the content of the published article.

## Acknowledgements

This work was supported by the National Science Foundation (NSF) under Award No. OIA-2418752 and NIH award No. P20GM103443. The authors gratefully acknowledge the contributions and support of Dr. Nick Klein and Dr. Dana Gehring (OLC), Dr. Badireddy (UVM), Dr. Tara Kulkarni (Norwich University), and Dr. Dubois (NMSU). Additional support was provided by the NSF 2D-BEST Center and the Department of Civil and Environmental Engineering at South Dakota Mines.

## Appendix A. Supplementary data

Supplementary data associated with this article are available in the Mendeley Data repository at: https://doi.org/10.17632/7b929fysdb

## Data Availability

The raw metagenomic sequencing data analyzed in this study are publicly available through the European Nucleotide Archive (ENA) under BioProject accession PRJEB68319. The corresponding ENA sample and sequencing run accession numbers for all wastewater metagenomic samples used in this study are provided in Supplementary Data 1. Environmental variables were retrieved from the NASA POWER database using the sampling coordinates and collection dates described in this study. The processed ARG feature matrix and supplementary datasets are available through Mendeley Data (Version 1, DOI: 10.17632/7b929fysdb.1). The Python source code used for data preprocessing, CatBoost model development, Leave-One-Group-Out (LOGO) cross-validation, 5-fold cross-validation, and SHAP analysis is publicly available at: https://github.com/bicbioeng/ARG_cross_city_prediction.

